# Post-tetanic potentiation lowers the energy barrier for synaptic vesicle fusion independently of Synaptotagmin-1

**DOI:** 10.1101/2020.08.14.251322

**Authors:** Vincent Huson, Marieke Meijer, Rien Dekker, Mirelle ter Veer, Marvin Ruiter, Jan van Weering, Matthijs Verhage, L. Niels Cornelisse

## Abstract

Previously, we showed that modulation of the energy barrier for synaptic vesicle fusion boosts release rates supralinearly (Schotten, 2015). Here we show that mouse hippocampal synapses employ this principle to trigger Ca^2+^-dependent vesicle release and post-tetanic potentiation (PTP). We assess energy barrier changes by fitting release kinetics in response to hypertonic sucrose. Mimicking activation of the C2A domain of the Ca^2+^-sensor Synaptotagmin-1 (Syt1), by adding a positive charge (Syt1^D232N^) or increasing its hydrophobicity (Syt1^4W^), lowers the energy barrier. Removing Syt1 or impairing its release inhibitory function (Syt1^9Pro^) increases spontaneous release without affecting the fusion barrier. Both phorbol esters and tetanic stimulation potentiate synaptic strength, and lower the energy barrier equally well in the presence and absence of Syt1. We propose a model where tetanic stimulation activates Syt1 dependent and independent mechanisms that lower the energy barrier independently in an additive manner to produce PTP by multiplication of release rates.

## Introduction

Synaptic transmission is a highly dynamic process. Vesicle release rates change several orders of magnitude in response to Ca^2+^ influx [1,2], and during repeated synaptic activity, the number of vesicles released by an action potential (AP) rapidly change [3]. Synaptic vesicle release is tightly controlled by specialized proteins, including SNAREs, SM proteins, and Ca^2+^-sensors, among others [4]. Many of these are involved in the last step of the release process in which the fusion of the lipid membranes of the vesicle and presynaptic terminal occurs. This process can be described in terms of an energy landscape. Here a fusion energy barrier represents the activation energy that is required for the intermediate steps during membrane fusion, such as overcoming electrostatic repulsion of the membranes [5] and lipid stalk formation [6]. In our previous work we showed that additive changes in the fusion energy barrier produced supralinear changes in the vesicle fusion rate, as predicted by transition state theory [7]. We hypothesized that an energy barrier model could explain the supralinear relationship between Ca^2+^-concentration and release rates [1] if Ca^2+^ binding to the release sensor would decrease the fusion energy barrier [7]. The latter assumption has not been tested experimentally. Synaptotagmin-1 (Syt1) is the Ca^2+^-sensor that is responsible for fast release in hippocampal synapses [8–10]. It was recently suggested that Syt1 lowers the fusion energy barrier by reducing electrostatic repulsion between membranes upon binding of Ca^2+^ [5]. However, Syt1 is also involved in translocating vesicles to the plasma membrane [11–14], and inhibiting spontaneous and asynchronous release [15–20]. Both these functions could potentially be involved in energy barrier modulation. Furthermore, in the absence of Syt1, slower Ca^2+^-sensors that drive asynchronous release become prominent [8,9,21]. Synaptotagmin-7 has been shown to trigger asynchronous release [22–24], but is also identified as a Ca^2+^ sensor for short-term facilitation [25,26]. The latter has been proposed to be due to its concerted action with Syt1 on the fusion energy barrier [7,25,27].

In this study, we tested the assumptions that (1) Syt1 can reduce the fusion energy barrier when activated, and (2) PTP can be produced by activation of a second pathway that reduces the fusion barrier independently of Syt1, thereby amplifying the action of Syt1. Changes in the energy barrier were probed in several mutant variants of Syt1 and during PTP, using hypertonic sucrose (HS) stimulation. We found that mimicking activation of Syt1’s Ca^2+^-binding C2A domain reduced the energy barrier. Syt1’s release inhibitory function acts independently from the energy barrier, indicating that additional release promoting factors contribute to spontaneous release. We found that after PTP or phorbol ester application the energy barrier was reduced, independently of the vesicle pool size and positional priming. Furthermore, this reduction did not require Syt1, and most likely is induced by activation of a second sensor. Altogether, these findings support a dual-sensor energy barrier model for supralinear Ca^2+^-sensitivity of release and changes in synaptic strength after PTP.

## Results

### Synaptotagmin-1 inhibits spontaneous release without changing the fusion energy barrier

Syt1 is well established to be the fast Ca^2+^-sensor for synaptic vesicle release in many synapses [8,10]. However, Syt1 is also know to act as an inhibitor of spontaneous and asynchronous release [15–20,28,29]. This inhibition may act directly on the fusion machinery [5,30,31], suggesting an increase in the fusion energy barrier [5]. Alternatively, Syt1 may inhibit a second high-affinity Ca^2+^-sensor [15,28], reducing sensitivity to local Ca^2+^-fluctuations [32–34], but likely not affecting the energy barrier. To discriminate between these two possibilities, we investigated whether the energy barrier in a resting synapse was altered in the absence of Syt1, comparing wild type (WT) and Syt1 KO glutamatergic hippocampal neurons. Syt1 deficiency did not affect the number of synapses (Fig. 1A,B) or dendrite length (Fig. 1A,C). However, electron microscopy (EM) revealed a significant reduction in membrane-proximal synaptic vesicles in Syt1 KO synapses (Fig. 1 Supp.1), as observed before [13]. Voltage clamp recordings revealed that spontaneous miniature excitatory post-synaptic current (mEPSC) frequency was more than doubled (Fig. 1E), while first evoked EPSC charge was strongly reduced in Syt1 KO synapses, compared with WT (Fig. 1F,G). As described previously, several synaptic parameters that provide information about priming and fusion [7] can be obtained by fitting a minimal vesicle state model to HS responses. These include the pool of primed vesicles or readily releasable vesicle pool (RRP) as defined by the total number of vesicles released by an osmotic shock from 500mM HS, maximal HS release rate (k_2,max_), and change in fusion energy barrier (ΔE_a_ (RT)) (For detailed methodology see: Schotten *et al.*, 2015). In order to examine changes in the energy barrier and the RRP, we applied a range of HS concentrations to WT and Syt1 KO synapses (Fig. 1H). As shown previously [7], kinetics of the responses became faster, while the delay of the onset of response decreased for increasing concentrations (Fig. 1 supp. 2). The readily releasable pool (RRP), quantified from model fits of the current response to 500mM HS, was not changed in Syt1 KO synapses (Fig. 1I), despite the reduced number of membrane-proximal vesicles found with EM. The fraction of the RRP depleted by 250mM sucrose (Fig. 1J), a proxy for the energy barrier height [5,35], was not changed. Beyond 250mM, HS-induced release rates even tended to be lower in Syt1 KO synapses than in WT (Fig. 1K), corresponding to an increased fusion energy barrier under these conditions (Fig. 1L). Therefore, we conclude that Syt1 in the non-activated state (i.e. without Ca^2+^ bound) does not increase the energy barrier for synaptic vesicle fusion, despite its inhibitory effect on spontaneous release.

**Figure 1.**
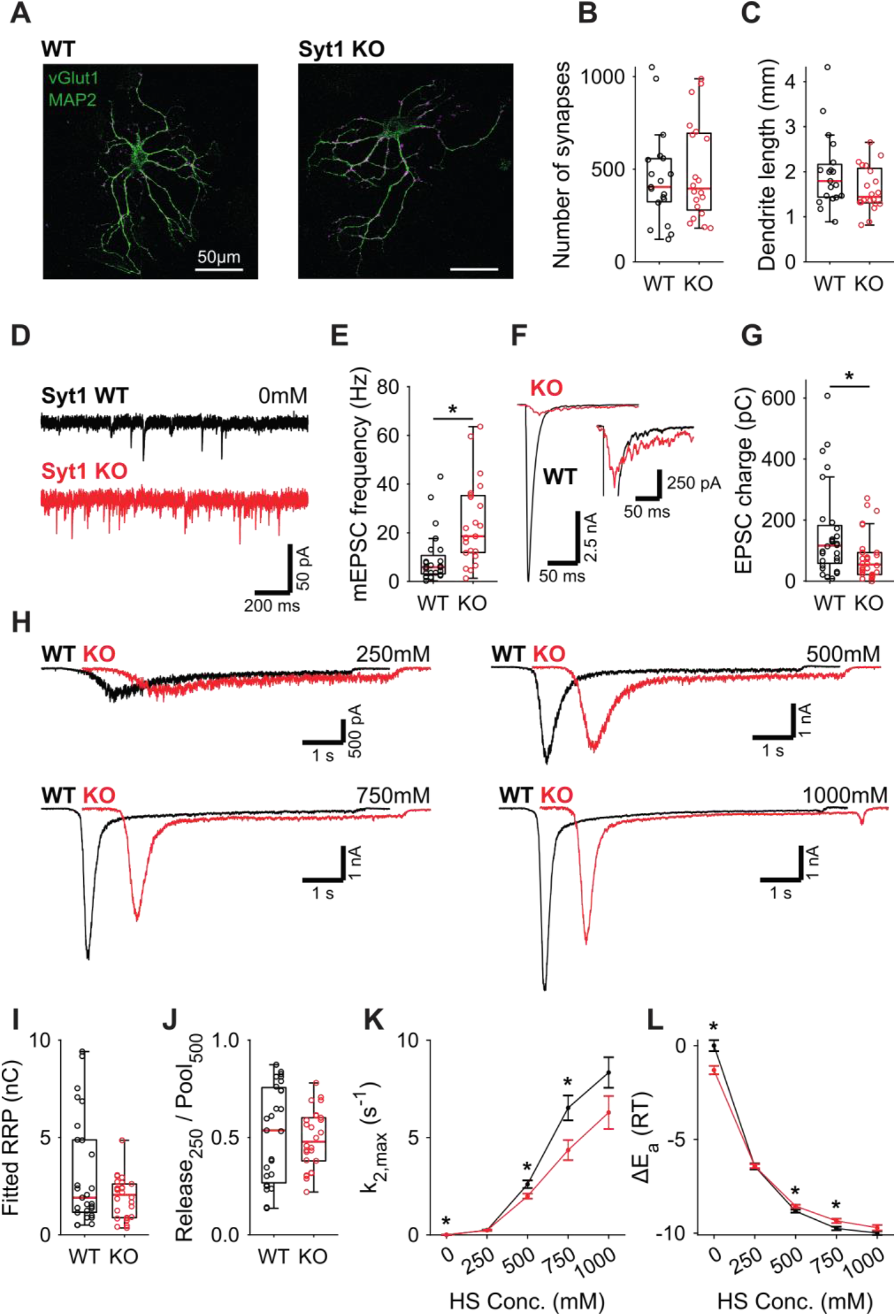
Increased spontaneous release in Syt1 KO does not correspond to a decrease in the fusion energy barrier. **(A)** Representative examples of WT and Syt1 KO neurons with vGlut (magenta) stained as a synapse marker and MAP2 (green) as a dendrite marker. **(B)** Boxplots of number of synapses, and **(C)** dendrite length per neuron in WT and Syt1 KO. **(D)** Representative traces of spontaneous release (0mM HS), and **(E)** boxplot of spontaneous frequency in WT and Syt1 KO. **(F)** Representative traces of AP evoked release in WT and Syt1 KO, overlaid, full view (left) and zoomed to the amplitude of Syt1 KO. **(G)** Boxplot of charge transferred during the first evoked EPSC. **(H)** Representative traces of HS induced release in WT and Syt1 KO overlaid with 1s offset, at 250mM, 500mM, 750mM, and 1000mM HS, and boxplots of **(I)** RRP charge estimated from 500mM HS, **(J)** depleted RRP fraction at 250mM HS in WT and Syt1 KO. **(K)** Plots (mean ± S.E.M.) of maximal HS release rates, and **(L)** change in the fusion energy barrier at different HS concentrations for WT and Syt1 KO. (* *p* < 0.05, Wilcoxon rank sum test).

### 9Pro linker mutant confirms inhibition of spontaneous release by Synaptotagmin-1 is not through an increase in the energy barrier

Syt1 KO abolishes synchronous release triggering [8–10] and impairs vesicle priming [11–14]. To control for potential effects from these properties on HS induced release, we used the Syt1 9Pro mutation to study the link between the inhibition of spontaneous release and the energy barrier in isolation. In this mutant, the flexible linker between Syt1’s two Ca^2+^-sensing C2 domains is fixed, selectively impairing inhibition of spontaneous release, without affecting evoked release [29,36]. To further minimize activation of Ca^2+^-sensors due to resting levels of intracellular Ca^2+^, all recordings, with the exception of first evoked release, were done in 0mM extracellular Ca^2+^ with 20μM BAPTA-AM. We expressed either Syt1 WT or Syt1 9Pro in Syt1 KO neurons, with Syt1 abundance in synapses exceeding endogenous levels to the same extent (Fig. 2 Supp. 1). We found a strong increase in mEPSC frequency in Syt1 9Pro synapses compared to Syt1 WT (Fig. 2A,B). The first evoked EPSC was unaffected (Fig. 2C,D). Applying 250mM and 500mM HS (Fig. 2E, Fig. 2 supp. 2), we found no difference in RRP (Fig. 2F), or depleted RRP fraction at 250mM (Fig. 2G). Release rates in Syt1 9Pro synapses in the presence of BAPTA-AM remained higher at 0mM sucrose (spontaneous release) compared with Syt1 WT (Fig. 2H). However, at 250mM and 500mM HS we found no effect on the release rates and corresponding fusion energy barriers (Fig. 2H,I). These data confirm that suppression of spontaneous release by Syt1 is not achieved by increasing the fusion energy barrier.

**Figure 2.**
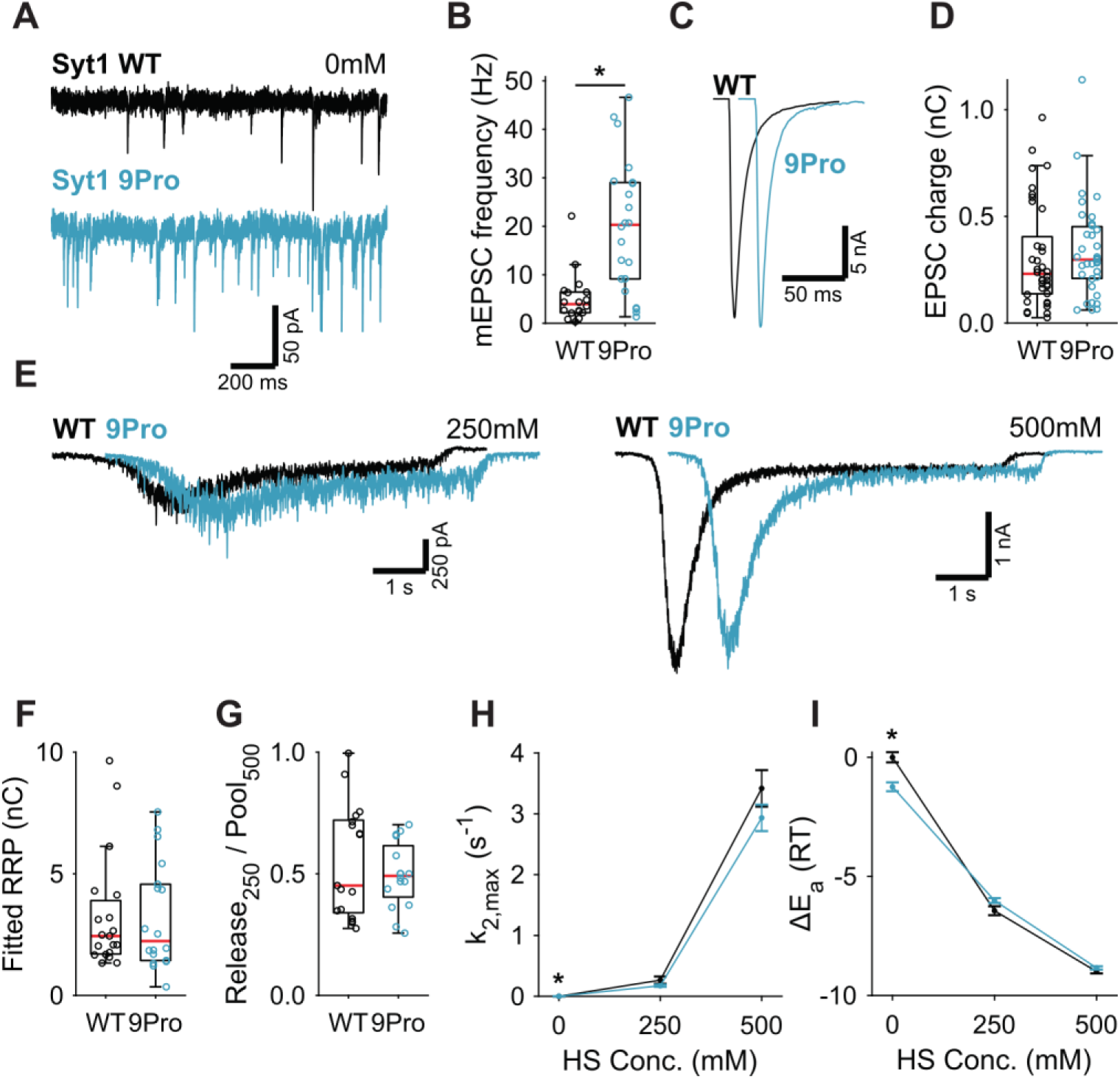
Syt1 9Pro mutation increases spontaneous release independent of the fusion energy barrier. **(A)** Representative traces of spontaneous release (0mM HS), and **(B)** boxplot of spontaneous frequency in Syt1 WT and Syt1 9Pro expressing synapses. **(C)** Representative traces of AP evoked release in Syt1 WT and Syt1 9Pro, overlaid with 20ms offset, and **(D)** boxplot of charge transferred during the first evoked EPSC. **(E)** Representative traces of HS induced release in Syt1 WT and Syt1 9Pro, overlaid with 1s offset, at 250mM and 500mM HS, and boxplots of **(F)** RRP charge estimated from 500mM HS, **(G)** depleted RRP fraction at 250mM HS in Syt1 WT and Syt1 9Pro. **(H)** Plots (mean ± S.E.M.) of maximal HS release rates, and **(I)** change in the fusion energy barrier at different HS concentrations for Syt1 WT and Syt1 9Pro. (* *p* < 0.05, Wilcoxon rank sum test).

### Activation of Synaptotagmin-1’s C2A domain lowers the fusion energy barrier

Next, we tested whether binding of Ca^2+^ to Syt1’s C2 domains lowers the energy barrier, as predicted by our energy barrier model for AP evoked release [7]. To be able to test this with our HS assay, which is too slow to detect energy barrier changes during AP induced Ca^2+^ binding, we analyzed two different Syt1 mutants that mimic the effect of persistent activation by Ca^2+^. We used the D232N mutation to neutralize a negatively charged residue in the C2A domain, which has been shown to increase Ca^2+^-triggered release [28,37]. Alternatively, we mimicked the Ca^2+^-mediated association with phospholipids by increasing the hydrophobicity of the C2 domains, through insertion of tryptophan mutations in the C2 domains [38] (M173W, F234W, V304W, I367W; Syt1 4W). Given the high peak currents observed in previous preparations, we recorded the Syt1 D232N gain-of-function mutant, and matching Syt1 WT, in 2mM extracellular Ca^2+^ to avoid voltage-clamp artifacts. Synaptic expression of Syt1 WT or Syt1 D232N in Syt1 KO neurons exceeded endogenous Syt1 levels to the same extent (Fig. 2 Supp. 1). We confirmed previous findings in mass cultures [28,37] of increased mEPSC frequency (Fig. 3A,B) and increased first evoked release in Syt1 D232N expressing synapses (Fig. 3C,D). HS evoked release revealed no significant difference in RRP size (Fig. 3E,F, Fig. 3 supp. 1A), but depleted RRP fraction at 250mM was increased in Syt1 D232N expressing synapses (Fig. 3G). Release rates at 175mM and 250mM HS were increased 1.7 to 1.9 fold, and the fusion energy barrier was reduced by 0.5 to 0.6 RT (Fig. 3H,I), while a similar trend was observed for higher concentrations. Hence, constitutively activating Syt1 by removing negative charge from its Ca^2+^-sensing domain increases vesicle fusion by decreasing the fusion energy barrier.

**Figure 3.**
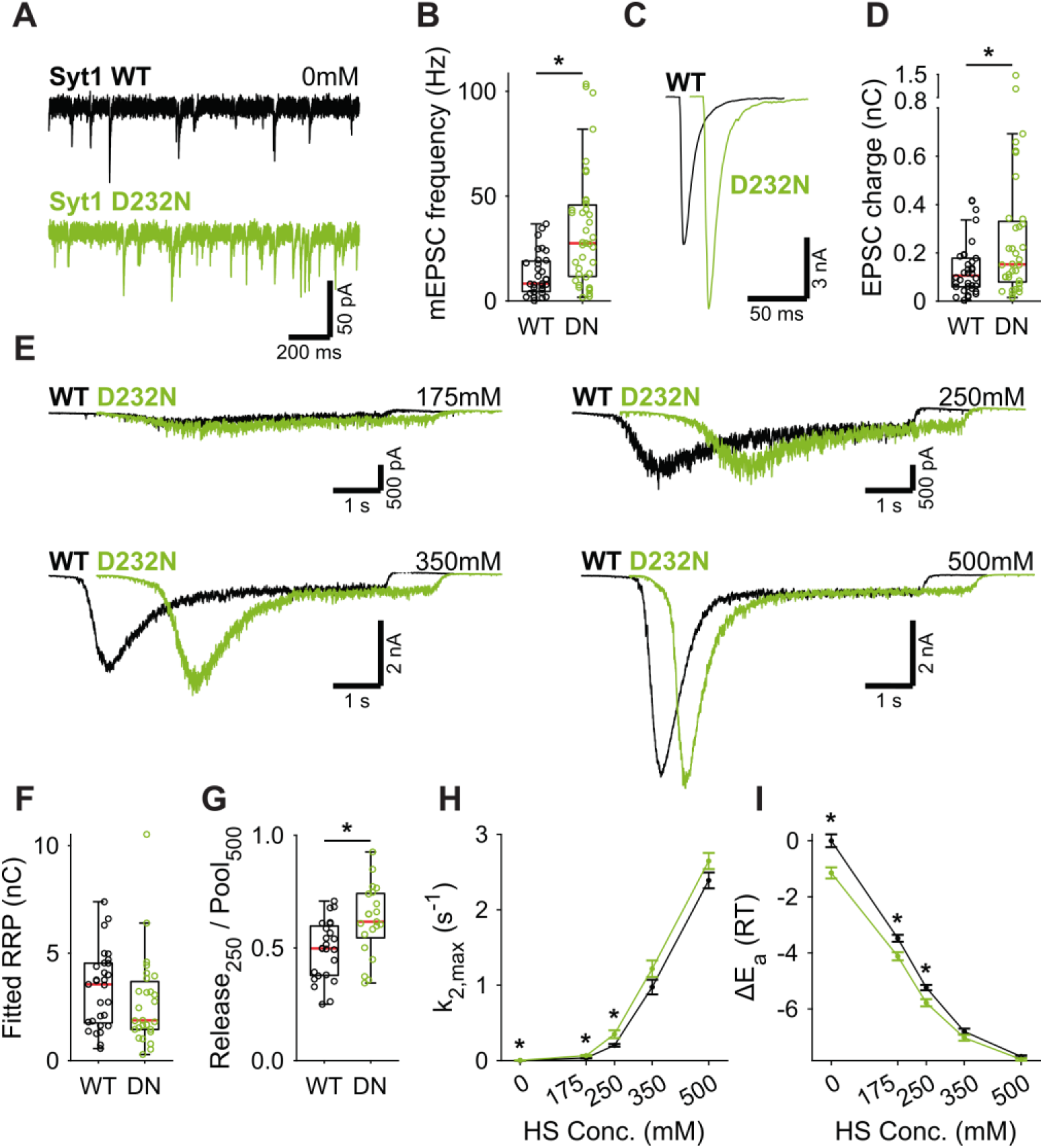
A decreased energy barrier increases vesicle fusion in Syt1 D232N expressing synapses. **(A)** Representative traces of spontaneous release (0mM HS), and **(B)** boxplot of spontaneous frequency in Syt1 WT and Syt1 D232N expressing synapses. **(C)** Representative traces of AP evoked release in Syt1 WT and Syt1 D232N, overlaid with 20ms offset, and **(D)** boxplot of charge transferred during the first evoked EPSC. **(E)** Representative traces of HS induced release in Syt1 WT and Syt1 D232N, overlaid with 1s offset, at 175mM, 250mM, 350mM, and 500mM HS, and boxplots of **(F)** RRP charge estimated from 500mM HS, **(G)** depleted RRP fraction at 250mM HS in Syt1 WT and Syt1 D232N. **(H)** Plots (mean ± S.E.M.) of maximal HS release rates, and **(I)** change in the fusion energy barrier at different HS concentrations for Syt1 WT and Syt1 D232N. (* *p* < 0.05, Wilcoxon rank sum test).

In Syt1 4W expressing synapses, we found a strong increase in mEPSC frequency (Fig. 3 Supp. 2A,B). No effect on first evoked charge was found (Fig. 3 Supp. 2C,D), but vesicular release probability was increased (Fig. 3 Supp. 2E). HS evoked release revealed no difference in RRP size (Fig. 3 Supp. 2F,G, Fig. 3 Supp. 3A). At 250mM HS the depleted RRP fraction (Fig. 3 Supp. 2H) and the release rate (Fig. 3 Supp. 2I) were increased, both indicating a reduction of the fusion energy barrier by 0.5 RT (Fig. 3 Supp. 2J). However, we found no significant differences at other HS concentrations, and at 500mM and 750mM release rates trended towards a decrease (Fig. 3 Supp. 2H). Such a reversal of phenotype cannot readily be explained through energy barrier modulation alone, suggesting an interaction between HS and the 4W mutations (see discussion). Our data at 250mM are in line with our findings with the Syt1 D232N mutant, corroborating the conclusion that activation of Syt1 C2A domain decreases the fusion energy barrier. Hence, supralinear Ca^2+^-sensitivity through multiplicative effects on the release rate [1,7] is likely supported by reductions of the energy barrier by Syt1.

### Induction of post-tetanic potentiation lowers the fusion energy barrier

Having established that activation of Syt1’s C2A domain reduces the fusion energy barrier, we next investigated whether induction of short-term plasticity (STP) through repetitive stimulation also leads to a reduction. We assessed changes in the fusion barrier after induction of PTP in the presence and absence of Syt1. PTP is a form of STP which lasts tens of seconds to minutes [39], allowing sufficient time to measure its effects using HS stimulation [40,41]. It has previously been suggested that PTP acts by decreasing the fusion energy barrier [40,41]. However, other studies have shown that an increase in the RRP after PTP can explain a large part of the potentiation of the EPSC [42,43]. To resolve this, we induced synaptic release either through APs or HS before (Naive; Fig. 4A), and 5 seconds after a train of 100 APs at 40Hz (PTP; Fig. 4A), to measure PTP, changes in the fusion energy barrier, and potentiation of RRP in the same cell. We have shown previously that our HS assay is sensitive enough to detect accelerated recovery of the RRP after 40Hz stimulation [44]. To maximize observable effects on the probability of vesicle release, we lowered extracellular Ca^2+^ to 1mM for this experiment. PTP increased EPSC charge by 39% (Fig. 4B,C), even though the RRP, as assessed from 500mM HS, was not fully recovered at this point (Fig. 4E,F, Fig. 4 Supp. 1A). This resulted in a 41% increase in vesicle release probability, calculated as the ratio of the AP and HS induced EPSC charges (Fig. 4D). PTP increased the depleted RRP fraction at 250mM (Fig. 4G), and increased HS release rates 17 to 37% (Fig. 4H), indicating about a 0.2-0.3 RT reduction of the fusion barrier (Fig. 4G,I). These results confirm previous findings that PTP does not increase the RRP [40,45], and shows that it is associated with a decrease in the fusion energy barrier.

**Figure 4.**
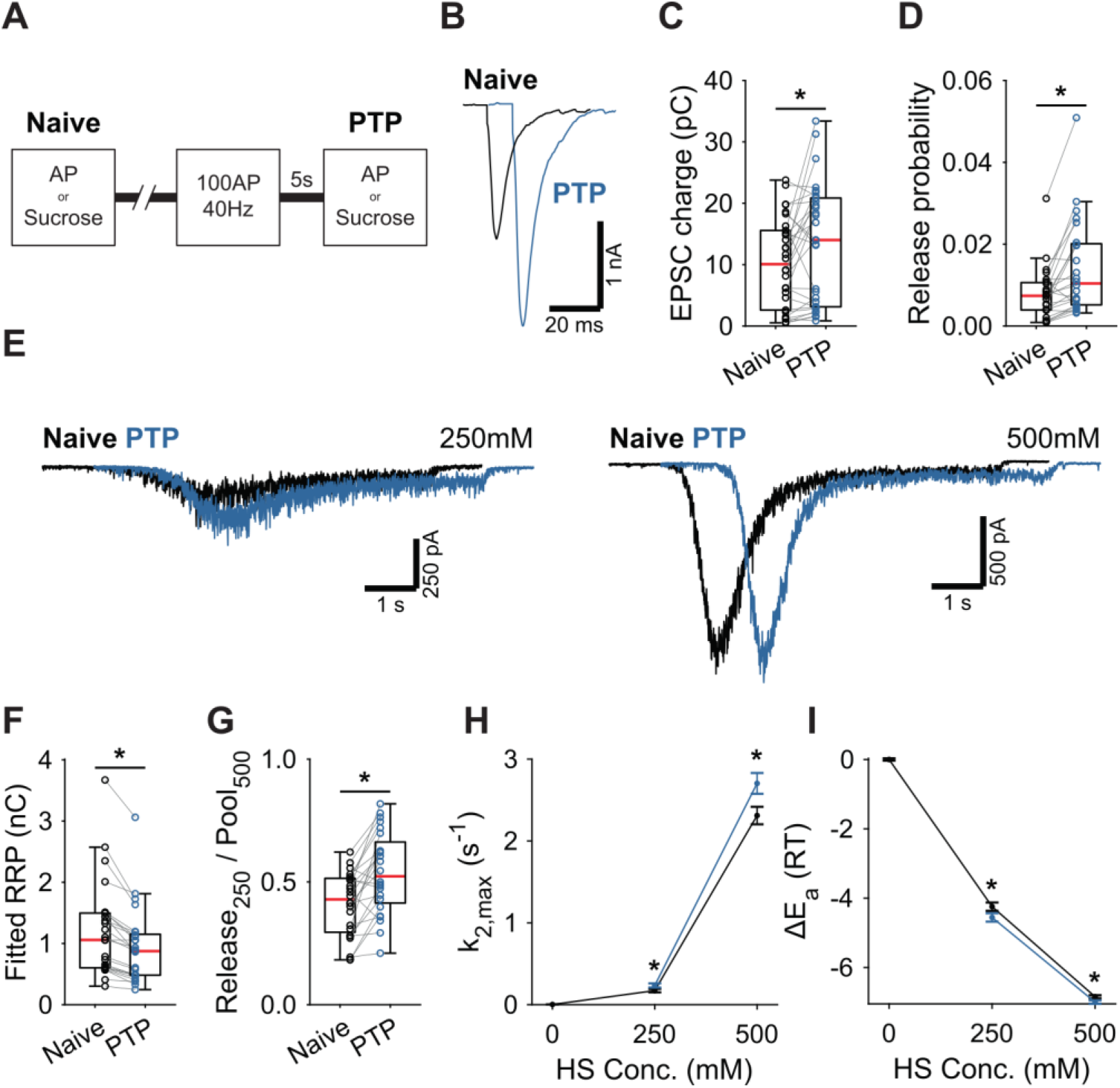
Post-tetanic potentiation causes a decrease in the fusion energy barrier. **(A)** Overview of protocol used to induce PTP. **(B)** Representative traces of AP evoked release before (Naive) and after PTP in WT synapses, overlaid with 10ms offset, and **(C)** boxplots of charge transferred during the first evoked EPSC and **(D)** release probability calculated by dividing the EPSC charge by the HS derived RRP charge **(F)**. **(E)** Representative traces of HS induced release before and after PTP, overlaid with 1s offset, at 250mM and 500mM HS, and boxplots of **(F)** RRP charge estimated from 500mM HS, and **(G)** depleted RRP fraction at 250mM HS before (Naive) and after PTP. **(H)** Plots (mean ± S.E.M.) of maximal HS release rates, and **(I)** change in the fusion energy barrier at different HS concentrations before (Naive) and after PTP. (* *p* < 0.05, Wilcoxon signed-rank test).

### PTP lowers the fusion energy barrier independently of Synaptotagmin-1

PTP induction is known to involve activation of Munc18-1 and PKC [43,46], but the role of synaptotagmin in this pathway is less clear. We showed previously that phosphorylation by PKC plays a role in Syt1-, but not in Syt2 expressing synapses [47]. However, whether Syt1 is required for the energy barrier reduction after PTP is not known. To investigate this, we performed a similar set of PTP experiments in Syt1 KO autapses, using 4mM extracellular Ca^2+^. In Syt1 KO autapses, PTP also increased vesicle release probability (Fig. 5A-C), while the RRP remained incompletely recovered (Fig. 5D,E, Fig. 5 Supp. 1A). Additionally, similar to WT autapses, we observed after PTP an increase in depleted RRP fraction at 250mM (Fig. 5F), and increased HS release rates (Fig. 5G), corresponding to a decrease in the fusion energy barrier of 0.3-0.6RT (Fig. 5H). Hence, reductions of the energy barrier due to PTP, do not require Syt1. Furthermore, as the RRP was not yet fully recovered after PTP, the contribution from the decreased energy barrier to potentiation of the EPSC is independent of priming. These findings suggest a model in which independent pathways for release cooperate to potentiate synaptic strength.

**Figure 5.**
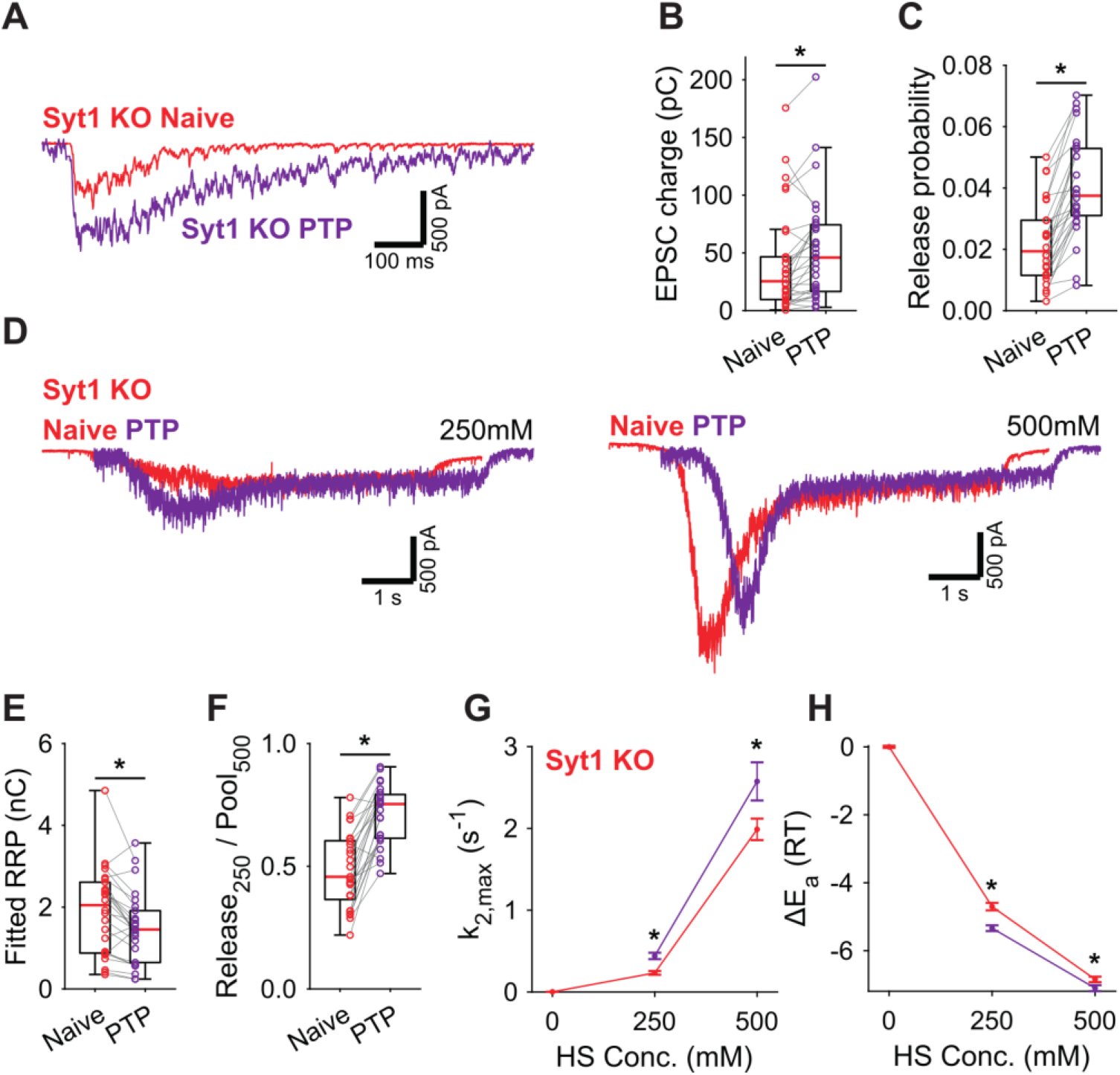
Post-tetanic potentiation decreases the fusion energy barrier independent of Syt1. **(A)** Representative traces of AP evoked release before (Naive) and after PTP in Syt1 KO synapses, overlaid, and **(B)** boxplots of charge transferred during the first evoked EPSC and **(C)** release probability calculated by dividing the EPSC charge by the HS derived RRP charge **(E)**. **(D)** Representative traces of HS induced release before and after PTP, overlaid with 1s offset, at 250mM and 500mM HS, and boxplots of **(E)** RRP charge estimated from 500mM HS, and **(F)** depleted RRP fraction at 250mM HS before (Naive) and after PTP. **(G)** Plots (mean ± S.E.M.) of maximal HS release rates, and **(H)** change in the fusion energy barrier at different HS concentrations before (Naive) and after PTP. (* *p* < 0.05, Wilcoxon signed-rank test).

### Activation of the diacylglycerol pathway lowers the fusion energy barrier independently of Synaptotagmin-1

PTP acts via the same pathway as phorbol ester mediated potentiation [35,46–50]. We showed previously that activation of the diacylglycerol (DAG) pathway with 1μM phorbol 12,13-dibutyrate (PDBu) reduced the fusion energy barrier [7]. In another study we showed that preventing Syt1 to be phosphorylated by PKC blocked potentiation of AP induced release by phorbol esters but not the potentiation of HS responses [47]. To gather further proof for a Syt1-independent pathway for energy barrier reduction after PTP we investigated whether the PDBu-induced reduction also occurred in the absence of Syt1. To this end, we compared the effect of 1μM PDBu on release in WT or Syt1 KO autapses. Spontaneous mEPSC frequency increased significantly in the presence of PDBu for both WT and Syt1 KO synapses (Fig. 6A-C). PDBu application did not affect the RRP assessed with 500mM HS (Fig. 6D,E), and induced a similar increase in depleted RRP fraction at 250mM, in WT and Syt1 KO synapses (Fig. 6F). Release rates increased after application of PDBu (Fig. 6G,H), associated with a similar decrease in the fusion energy barrier (Fig. 6I,J) for both WT (0.4-0.6RT) and Syt1 KO (0.4-0.6RT). We conclude that both tetanic stimulation and activation of the DAG pathway reduces the fusion energy barrier, independently of Syt1. This supports a model for Ca^2+^-dependent release and PTP where release sensors are activated independently, but through additive effects on the fusion energy barrier produce multiplicative effects on the fusion rate.

**Figure 6.**
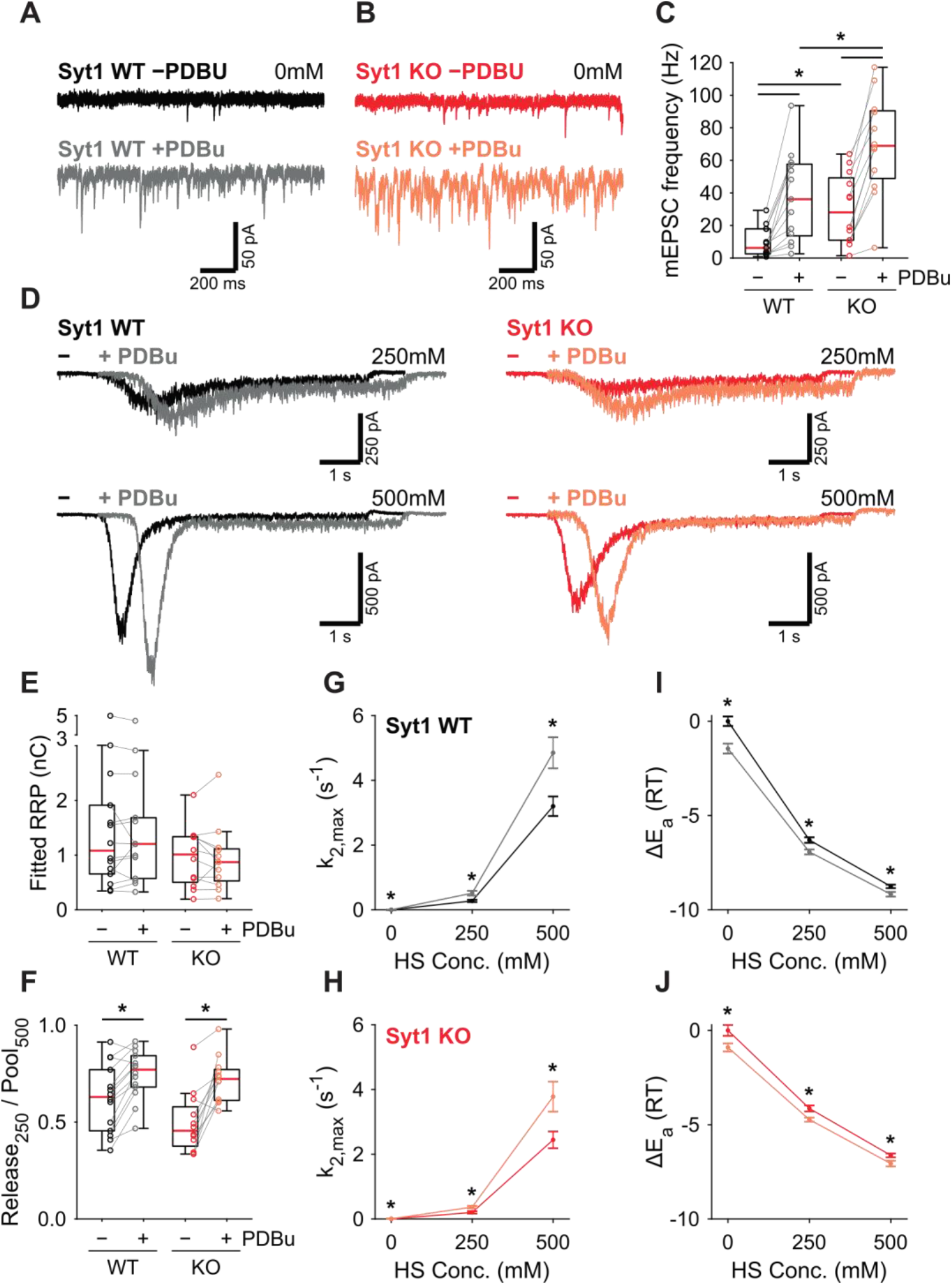
PDBu increases vesicles fusion by lowering the energy barrier in the presence or absence of Syt1. Representative traces of spontaneous release (0mM HS) before (−PDBu; top) and after (+PDBu; bottom bath application of PDBu in WT **(A)** and Syt1 KO **(B)** synapses. **(C)** Boxplots of spontaneous release frequency in WT and Syt1 KO before and after PDBu. **(D)** Representative traces of HS induced release at 250mM and 500mM HS, and boxplots of **(E)** RRP charge estimated from 500mM HS, **(F)** depleted RRP fraction at 250mM HS in WT and Syt1 KO before and after PDBu. **(G,H)** Plots (mean ± S.E.M.) of maximal HS release rates, and **(I,J)** change in the fusion energy barrier at different HS concentrations before and after PDBu, for WT **(G,I)** and Syt1 KO **(H,J)**. (* *p* < 0.05, Wilcoxon rank sum test for independent and Wilcoxon signed-rank test for paired samples).

### An energy barrier model for Ca^2+^ induced vesicle release and post-tetanic potentiation

Previously, we proposed that transition state theory can be used to describe the process of vesicle fusion. In this framework, the fusion reaction is defined as the transition from the primed state to the fused state, with *E*_*a*_ the activation energy that is required for the transition to occur, also referred to as the fusion energy barrier (Fig. 7A) [7]. According to the Arrhenius equation, there is an exponential relation between the activation energy and the rate of a reaction. This, implies that additive changes in the height of the fusion energy barrier lead to multiplicative effects on the fusion rate [7]. We showed that supralinear Ca^2+^-dependence of release follows from this principle, when assuming an energy barrier reduction Δ*E*_*f*_ for each Ca ion that binds (Fig. 7C, upper row) [1,7]. We now propose that the same framework can be used to describe asynchronous release and STP. This can be realized by adding additional release sensors to the model and adding their effects in the energy barrier domain (Fig. 7B) (Schotten *et al.*, 2015; eq. 4). When multiple sensors are activated (Fig. 7A,B), the new fusion energy barrier height *E*_*new*_ is given by:

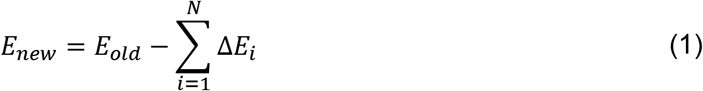

with *N* the number of sensors, Δ*E*_*i*_ the energy barrier reduction induced by sensor *i*, and *E*_*old*_, the basal energy barrier height. Applying the Arrhenius equation [7] gives the corresponding new fusion rate *k*_*2,new*_:

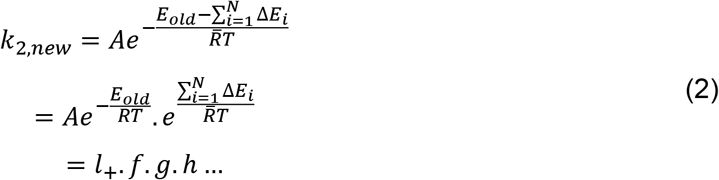

with 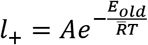 the basal fusion rate, 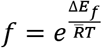 the factor by which *l*_+_ needs to be multiplied to account for activation of sensor *f*, and each additional sensor (i.e. *g*,*h*,…) after that (Fig. 7B). *A* is an empirical prefactor, 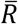 the gas constant, and *T* the temperature. During synaptic activity these sensors may be directly activated by Ca^2+^, or indirectly, through other pathways such as the diacylglycerol (DAG) pathway [51]. Through differences in activation and kinetic properties of different sensors, their combined effect could give rise to a diverse repertoire of vesicle release patterns in response to different patterns of presynaptic activity.

**Figure 7.**
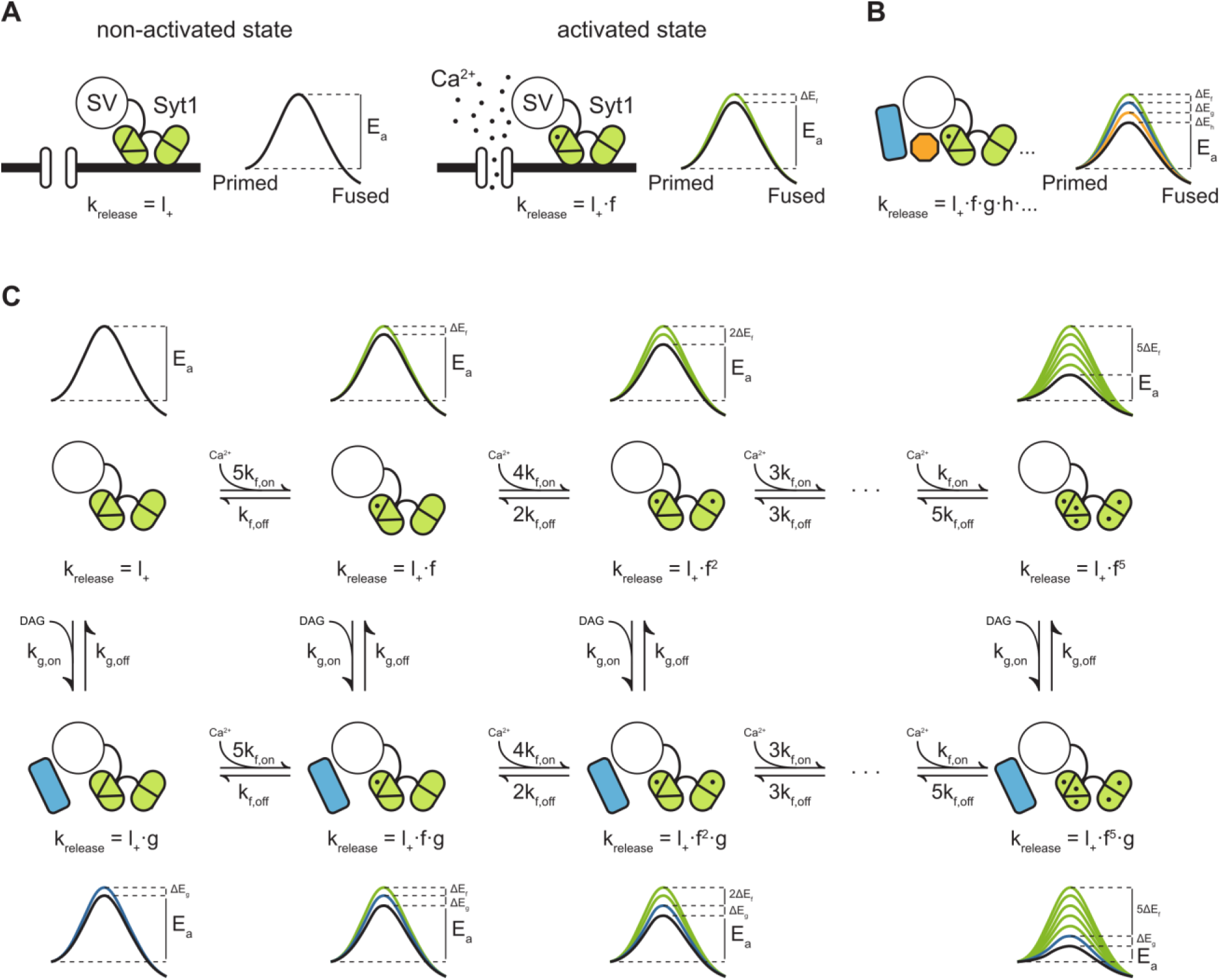
Modulation of the energy barrier through multiple sensors and activation states. **(A)** Fusion sensors in the non-activated state do not affect the fusion energy barrier (*E*_*a*_), release rates (k_release_) correspond to the energy barrier in the ground state (*l*_+_; left). Upon activation by Ca^2+^, Syt1 lowers the fusion energy barrier (Δ*E*_*f*_), multiplying release rates (*f*;right). **(B)** Multiple sensors in the activated state, each with separate additive effects on the energy barrier, provide multiplicative effects on release rates. **(C)** Binding of multiple Ca^2+^ ions to Syt1 up to a maximum of 5, may be represented as multiple activation states additively lowering the energy barrier and multiplying release rates (top row). Additional activation of a second sensor (*g*) further expands the total number of states (bottom row), providing additional release pathways and increasing potential for plasticity.

Based on our experimental findings, we propose a qualitative model for PTP, by combining the fast Ca^2+^ sensor with five activation states from the allosteric model [1,7], with a slow DAG sensor with one activation state. This gives a two-dimensional reaction scheme with in total 12 different vesicle states, each with its own associated fusion barrier (Fig. 7C). As in the original allosteric model [1], substantial activation of the fast sensor only occurs by peak Ca^2+^ levels reached during the AP due to its low Ca^2+^-affinity, which synchronizes vesicle fusion to the moment of AP firing (Fig.7C; upper row). Activation of the slow sensor can occur by increased DAG levels triggered by elevated residual Ca^2+^ after synaptic activity. The reduction of the fusion energy barrier Δ*E*_*g*_ after activation of the sensor multiplies the spontaneous release rate by a factor 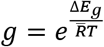 (Fig. 1C; left column), leading to asynchronous release. PTP is induced when both sensors are activated simultaneously, for instance when several seconds after the train stimulation a new AP triggers the activation of the fast sensor, while the slow sensor is still activated by residual DAG. At this point fusion rates of the fast sensor are multiplied with the multiplication factor *g* of the slow sensor (Fig. 7C; bottom row). Such a model provides a simple explanation of how additional release promoters with small individual effects, may achieve meaningful changes in EPSC size [7,25,27]. We explored how the Syt1 D232N mutation could affect Ca^2+^ sensitivity in this model. Based on proposed electrostatic effects on the fusion barrier [5], the effect of adding one charge to the C2A domain can be modelled by increasing the basal fusion rate *l*_+_ to 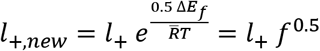. This is under the assumption that one charge is half as effective as two charges during Ca^2+^ binding [5], with Δ*E*_*f*_ the energy barrier reduction from one bound Ca^2+^ (Fig. 7. Supp. 1A,B). A two-fold increase in SNARE binding was also reported for this mutation, though only after Ca^2+^-binding [37]. Therefore, this does not affect the basal release rate *l*_+_, but can be modelled by replacing *f* with *f*_*D232N*_ = 2*f* (Fig. 7. Supp. 1A,B). Both parameter changes result in an increase in Ca^2+^-sensitivity of release, with higher release rates at all Ca^2+^ concentrations, and a similar small reduction in cooperativity (Fig. 7. Supp. 1B), while spontaneous release was only increased with increased *l*_+_.

## Discussion

Fusion rates change exponentially in response to linear changes of the fusion energy barrier. This makes modulation of the energy barrier a powerful principle for inducing quick and sizeable changes in synaptic strength. In this work, we propose that independent modulation of the energy barrier by different sensors in the synapse contributes to the supralinearity of Ca^2+^-dependent release and STP. In line with this idea, our results indicate that activation of the Ca^2+^-binding C2A domain of Syt1 potentiates release and decreases the energy barrier. Additionally, tetanic stimulation and PDBu both increase synaptic strength and decrease the fusion energy barrier, independent of Syt1. We propose that combined energy barrier reductions by Syt1 and the DAG-pathway contribute to the potentiation of EPSCs after PTP.

### Changes in spontaneous release do not directly correspond to changes in the energy barrier as assessed with hypertonic sucrose

When all sensors are in the non-activated state, the basal release rate constant *l*_+_ in our model is associated with one effective energy barrier representing the fusion pathway. This includes intermediate steps such as stalk formation, hemifusion and pore formation (Fig. 7A) [1]. In other studies its height has been estimated to be 30 k_B_T based on experiments with pure lipid bilayers and from course-grained simulations of the underlying intermediate fusion states [52–54]. Previously, we concluded that changes in spontaneous release rates after genetic or biochemical manipulations did not correspond with the energy barrier shifts measured with HS [7]. Here we showed both in Syt1 KO (Fig. 1) and Syt1 9Pro expressing synapses (Fig. 2) that mEPSC frequencies were increased, in line with previous studies in autapses [5,20,55] and networks [28,29,36,56]. It remains enigmatic why others found a similar increase in spontaneous release in networks but not in autapses after Syt1 deletion [57,58], but differences in culture protocol [59], or genetic background could play a role. Despite the increase in spontaneous release in our Syt1 KO and Syt1 9Pro expressing synapses, the fusion energy barrier assessed with HS was not changed (Fig. 1L; Fig. 2I). These findings indicate that for hippocampal autapses the rate constant for spontaneous release is not equal to the basal release rate constant *l*_+_ as assessed with HS in the model, and that additional mechanisms may also contribute to AP independent release of vesicles. As spontaneous release is to a large extend Ca^2+^ dependent [28,60], a possible mechanism may be rapid spontaneous Ca^2+^ fluctuations [32–34]. When such fluctuations occur locally at individual fusion sites, Ca^2+^ activation of a release sensor will reduce the energy barrier only locally and very briefly and not constitutively across all synapses simultaneously. We estimated previously that in such a scenario the frequency of Ca^2+^ fluctuations increases the release rate constant in an additive manner by 2-4 10^−4^ s^−1^ in WT autapses [7]. This will dominate the release rate at 0mM sucrose but is negligible compared to release rates induced by higher concentrations, corresponding to undetectable changes in the energy barrier. Interestingly, recording spontaneous release in 0mM extracellular Ca^2+^ and in the presence of the 20μM BAPTA seemed to reduce its frequency less drastically as reported in other studies (compare Fig. 1E and 2B) [28,56]. Moreover, differences in mEPSC frequency in Syt1 9Pro expressing synapses remained under these conditions. This might suggest that in our experiments the BAPTA loading was insufficient to block all spontaneous events. BAPTA-AM was applied after recording evoked release, with its incubation time restricted to 10 min to ensure the recording remained stable enough for HS measurements. Alternatively, part of the spontaneous release could be Ca^2+^ independent and through a different pathway than evoked release, either from a subset of synapses [61,62] or from a different vesicle pool [63,64]. In this case, changes in the fusion barrier of this pathway could have been missed if spontaneous released vesicles are a small subset of the total pool, or when these vesicles are somehow less sensitive to HS stimulation.

### Syt1 does not inhibit spontaneous release by increasing the fusion energy barrier as assessed with hypertonic sucrose

Several mechanisms have been proposed to explain the inhibitory effect of Syt1 on spontaneous release [17,18,28,29]. These include Syt1 clamping a second sensor for slow release [15,21], clamping fusion directly by arresting SNARE complexes [30,31], or increasing electrostatic repulsion between lipid membranes [5]. The latter two mechanisms imply an increase in the energy barrier in the presence of Syt1, which we did not observe with our HS assay (Fig. 1). Furthermore, Syt1 9Pro expressing synapses showed increased spontaneous release, but no changes in the energy barrier and normal AP induced release. This indicates that Syt1’s inhibitory role on spontaneous release is independent of its release-promoting role. We conclude that Syt1 does not increase the fusion energy barrier in its non-activated state. This conclusion does not support a model in which Syt1 suppresses spontaneous release by inhibiting the fusion step itself. Our data is most consistent with either a model where Syt1 clamps a slow sensor with high affinity for Ca^2+^, making the system less sensitive to spontaneous Ca^2+^ fluctuations, or with an additional fusion pathway for spontaneous release for a subset of vesicles, under the control of the clamping function of Syt1 (see discussion above).

### Activation of Syt1’s Ca^2+^ binding domain decreases the fusion energy barrier

A central assumption of our model is that activation of Syt1 by Ca^2+^ binding lowers the fusion barrier [5,27,65,66]. Several competing models have been proposed for this [66]. Both the Syt1 D232N mutant, where Ca^2+^-binding is mimicked, and the Syt1 4W mutant where hydrophobicity is increased, showed increased spontaneous and HS-induced release rates up to 250mM HS, in line with a reduced fusion energy barrier. Interestingly, for 500mM and 750mM we also found a slightly reduced fusion barrier in WT cells compared to Syt1 KO cells (Fig. 1L), possibly due to activation of Syt1 at basal Ca^2+^ levels. An electrostatic energy barrier model has been proposed, which assumes Ca^2+^-binding to Syt1 reduces the fusion barrier by diminishing electrostatic forces between opposing membranes [5]. Adding a single positive charge to the C2A domain in the D232N mutant yielded a reduction between 0.5 and 0.6 RT. Assuming linear scaling of the fusion barrier with charge [5], this implies a 5 to 6 RT (or 2.9 to 3.5 kCal mol^−1^) reduction when Syt1 is fully activated after binding 5 Ca^2+^ ions, adding 10 charges in total [1]. This is in the same order of magnitude as the estimated 10 RT (5.9 kCal mol^−1^) reduction during AP induced release in hippocampal neurons [38], but about 3 times smaller than the 17 RT reduction predicted for full occupation of Synaptotagmin 2 (Syt2) in the calyx of Held [1]. These results suggests that either differences exist in the efficiency of the fusion machinery in the two systems, or that our method has a limited ability to capture the full effect of Syt1 activation on the fusion barrier (see also discussion below). Alternatively, the energy barrier reduction and increased spontaneous release rate in the D232N mutant do not come from electrostatic effects, but from increased Ca^2+^-dependent SNARE binding [37] at basal Ca^2+^ levels. Modeling these scenario’s (reduced electrostatic repulsion or increased Ca^2+^-dependent SNARE binding) with the energy barrier model, predicted in both cases an increase in Ca^2+^ sensitivity with little effect on cooperativity (Fig. 7 supp. 1). These results are in line with previous studies in GABAergic neurons [28,37] with similar values for the apparent cooperativity. However, caution has to be taken to directly compare model predictions to these experimental studies, which show the IPSC amplitude (not peak release rate) as a function of the extracellular (not intracellular) Ca^2+^concentration. Interestingly, adding a positive charge to the C2A domain at a different position (D238N) was shown to reduce Ca^2+^ sensitivity in the same study [37]. This makes increased Ca^2+^-dependent SNARE binding a more plausible explanation for fusion barrier effects in the D232N mutant, but more research is needed.

In contrast to our HS measurements after PDBu [7] or PTP stimulation, we could not detect changes in the fusion barrier with concentrations beyond 250mM for these Syt1 mutants. This may point to a possible interaction of the mechanisms by which sucrose and Syt1 induce vesicle fusion, which may be a limitation of the method for some specific conditions. Properties that contribute to the lateral pressure of the membrane, such as membrane fluidity, bilayer thickness, hydration state of lipid headgroups, and interfacial polarity and charge, can change in response to osmotic pressure [67]. This could render the membrane bending properties of Syt1 less effective or different at higher sucrose concentrations. In case of the Syt1 4W mutation, opposite effects of increased hydrophobicity of the C2 domains on membrane fusion could dominate at different osmotic conditions. Enhanced affinity for phospholipids may promote membrane-membrane interactions, increasing the chance to cross the fusion barrier during no or mild osmotic stress. At conditions where the energy barrier is substantially reduced, deeper insertions of the C2AB domains in the plasma membrane may lead to less curvature, rendering the vesicles less fusogenic [68]. Based on these considerations, we conclude that activation of Syt1 by Ca^2+^ promotes vesicle release by reducing the fusion energy barrier.

### Energy barrier modulation contributes to PTP independently from vesicle priming

PTP and activation of the DAG-pathway are well established to increase synaptic strength [39,51]. Previous studies have suggested that energy barrier modulation contributes to STP [7,25,27,35,40,41], while other studies suggested increased vesicle priming [42,43] or decreased un-priming [69]. Using our HS assay, we found a decreased fusion barrier, but incomplete recovery of the RRP, after PTP (Fig. 4, 5). Furthermore, we found no change in RRP size after application of PDBU (Fig. 6E), in line with previous studies [7,35,46,70], except one [71]. This indicates that, in cultured hippocampal neurons, lowering of the fusion barrier, and not increased vesicle priming or decreased un-priming, is a major factor in PTP. This is in line with the notion that activation of the DAG-pathway only increases the “effective” pool size of AP releasable vesicles [70,72,73]. This process, also referred to as ‘superpriming’, involves conversion of slowly releasing vesicles into rapidly releasing vesicles [74–78], and could be interpreted as a transition of vesicles from a high-to low energy barrier state in the same RRP. PTP and PDBU stimulation reduced the fusion barrier about 0.2-0.6 RT and 0.4-0.6 RT, respectively (Fig. 4–6). Although these effects are small compared to the estimated 30 RT fusion barrier for pure lipid bilayers [52] they correspond to a multiplication of the fusion rate by a factor 1.2-1.8 and 1.5-1.8 (Eq. 2). These values are close to the 1.4 and 1.5-1.9 fold increase in evoked release after PTP (Fig. 4) and stimulation of the DAG pathway [35,46,50] in hippocampal autapses, and suggest a large contribution of fusion barrier modulation to tetanic-stimulation induced STP.

### PTP lowers the fusion energy barrier independently of Syt1

We postulated that PTP occurs through activation of a pathway that lowers the energy barrier independently of Syt1. Indeed, we found a similar reduction of the energy barrier after PDBu application or PTP induction in the presence or absence of Syt1. This reduction most likely requires activation of a second sensor [15,21]. According to our model, a second sensor both amplifies the Syt1 induced fusion rates and triggers release by itself (Fig. 7). However, the latter may occur at a slower rate than Syt1 induced release, and represent the increase of asynchronous release after tetanic stimulation [19,20] and the increase in spontaneous release after PDBu (Fig. 6). For PTP different sensors and/or pathways may be involved [39,51], including the DAG pathway. DAG-analogs enhance spontaneous release, AP induced release, and vesicle priming and reduce the fusion energy barrier [1,7,35,42,43,46–48,70,79]. Munc13 is directly activated by DAG [80], contributes to short-term synaptic plasticity [35,50,81–84] and modulates the fusion barrier [35]. PKCs have been identified as relevant DAG targets in hippocampal neurons [46] and the Calyx of Held [43]. Phosphorylation of synaptotagmin by PKC has been shown to play a role in STP in Syt1-, but not Syt2-, expressing synapses [47]. Interestingly, preventing phosphorylation of Syt1 did not affect the fusion barrier and an effect on priming was proposed.

In conclusion, lowering of the fusion energy barrier during PTP occurs independently of Syt1, likely by Ca^2+^ and/or DAG activation of an additional, slow sensor. However, an alternative mechanism is a reduction of electrostatic repulsion between opposing negative membranes through accumulation of residual Ca^2+^ [5,85].

### Fusion energy barrier model for PTP

Several mechanisms have been proposed for synaptic plasticity, including ways to increase the presynaptic Ca^2+^ signal (for review see [27]), (activity dependent) channel-attachment of vesicles [86,87], Ca^2+^ dependent vesicle priming [42,43] or inhibition of un-priming [69], and tightening of the SNARE-complex [78]. Our model employs a single mechanistic principle, multiplicative modulation of fusion rates through additive modulation of the energy barrier, to describe supralinear Ca^2+^-sensitivity of release and PTP. A different dual sensor model was proposed before in which release can only occur from the vesicle states where one or both of the sensors are fully activated by Ca^2+^ [21]. This model can accurately describe Ca^2+^-sensitivity of release in Syt1 WT and KO synapses but not STP. In our model, release can occur from all states, also when sensors are partly activated. This provides a general framework for STP in which multiple sensors can cooperate in controlling release through their additive effect on the fusion barrier, without requiring direct interaction. Within this framework, neurons can implement different forms of STP within the same synapses, by expressing different combinations of sensors, all with their own dynamic properties [51]. In addition, it can reconcile the dual role of slow sensors in release, being both a sensor for asynchronous release and STP. For paired-pulse facilitation, a form of STP occurring at a shorter time scale than PTP [51], such a dual role has been suggested for Syt7 in corticothalamic and hippocampal synapses [22–27,88]. Its role as a facilitation sensor, with a multiplicative effect on the fusion rate, is supported by the finding that multiplication of membrane bound Syt1 and Syt7, as a proxy for the activation of both sensors, can account for the observed facilitation [27]. However, a dual sensor model could not explain all features of vesicle release in the Drosophila neuromuscular junction [69]. In its current form our model for PTP is a qualitative and reduced model which does not include (Ca^2+^-dependent) priming of vesicles, spatial distribution of vesicles or detailed Ca^2+^ dynamics, nor the effect of Syt1 or other sensors on these processes. Furthermore, it does not explicitly describe all successive steps in the fusion pathway [6] and the different roles of Syt1 during these steps [14,89,90], but models fusion as a single step using one effective energy barrier. Despite these limitations, the model is able to explain the increase in fusion rate upon Ca^2+^ binding and increase in release probability during PTP, the latter with the relatively small changes in fusion barrier we found. This suggest that, while other mechanisms also may contribute, modulation of the fusion barrier is an important mechanism by which the synapse controls its efficacy on the short-term scale.

## Materials and Methods

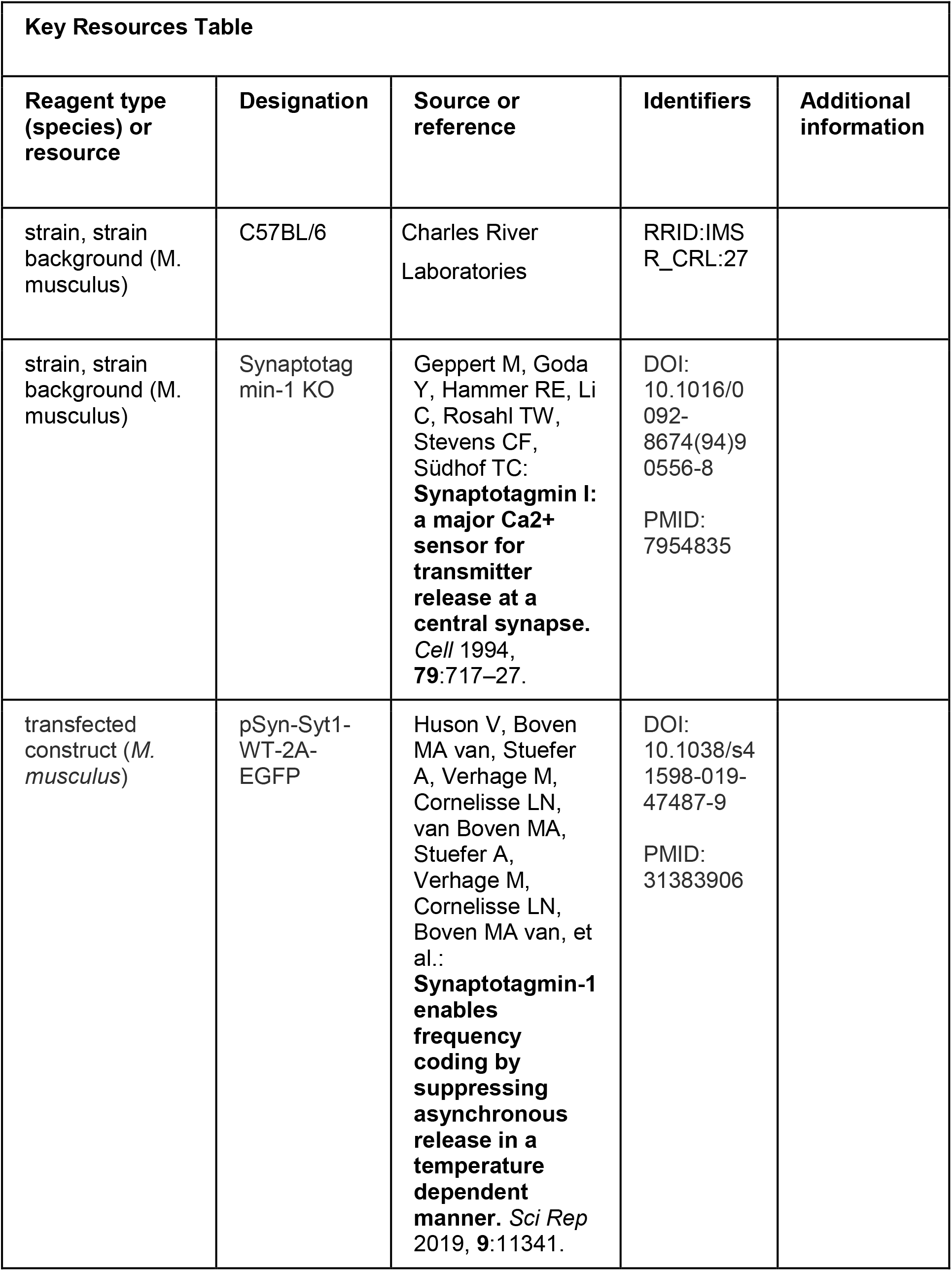
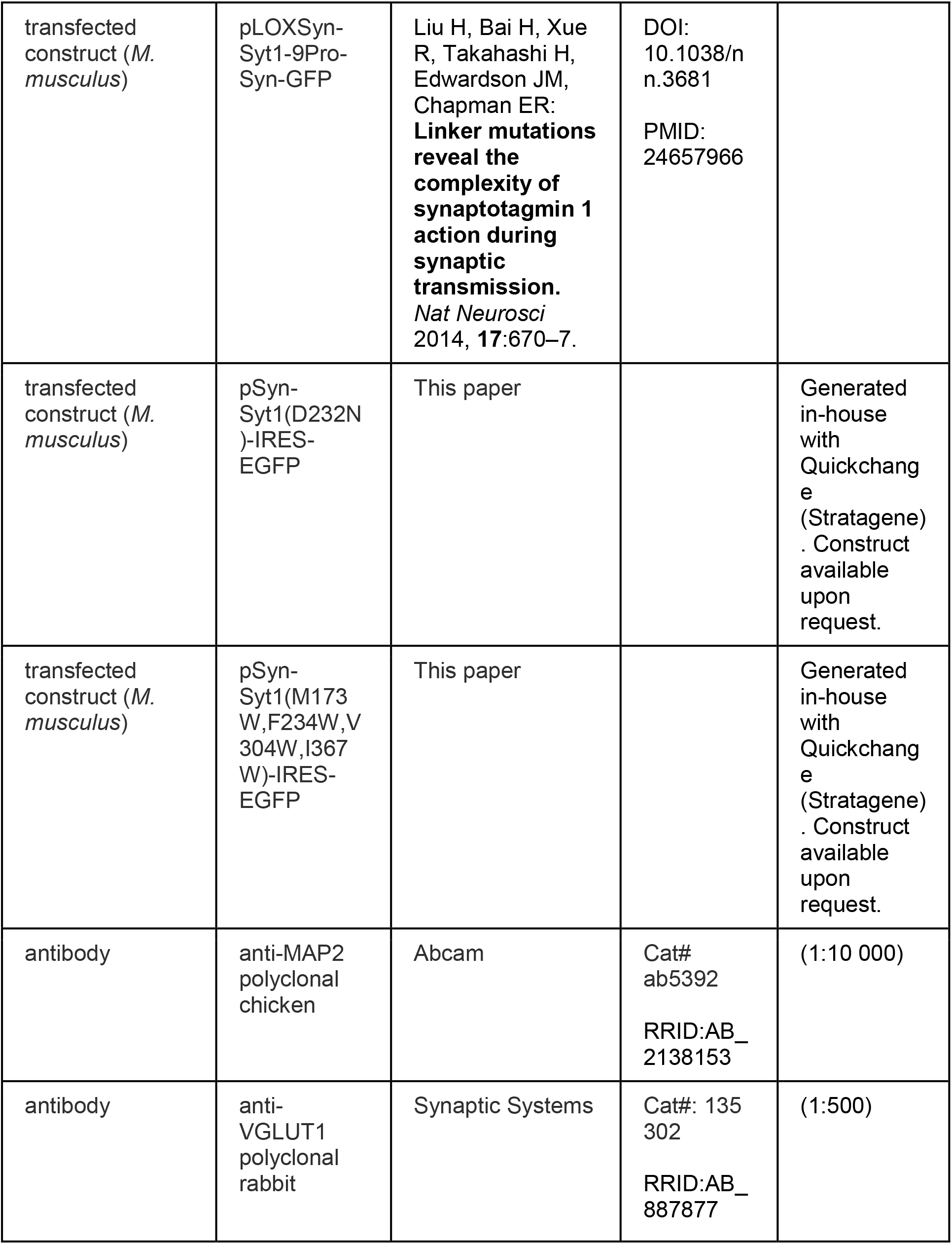
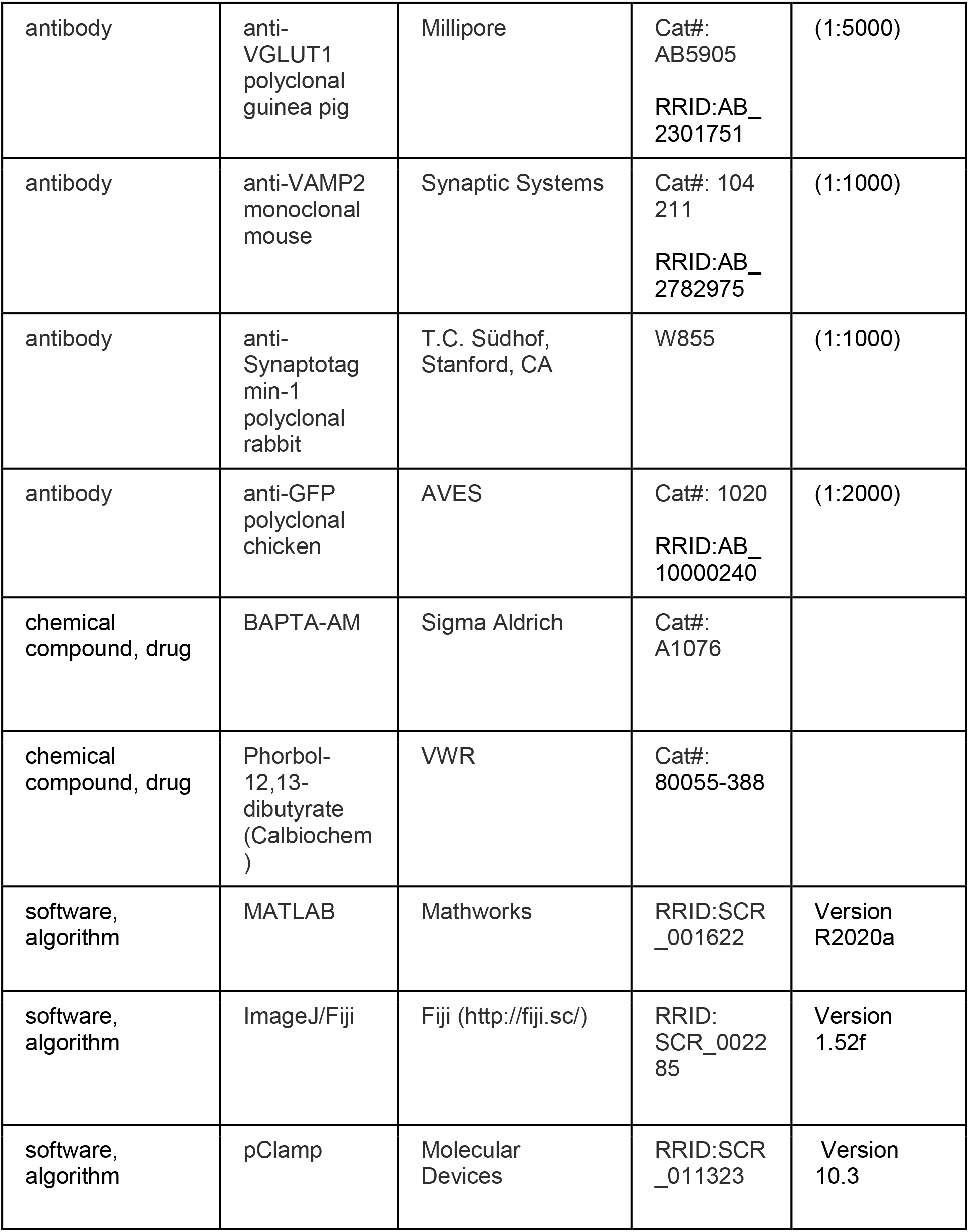

### Animals

Neuronal cultures were prepared from embryonic day 18 (E18) pups of both sexes, obtained by caesarean section of pregnant female mice. For this, previously described Synaptotagmin-1 knockout [8] or C57BL/6 mouse lines were used. Newborn pups (P0-P1) from Winstar rats were used for glia preparations. Animals were housed and bred according to institutional and Dutch governmental guidelines, and all procedures are approved by the ethical committee of the Vrije Universiteit, Amsterdam, The Netherlands.

### Dissociated Neuronal Cultures and Lentiviral Infection

Hippocampi from WT and Syt1 KO mice were isolated, collected in ice-cold Hank’s buffered salt solution (HBSS; Sigma) buffered with 1mM HEPES (Invitrogen), and digested for 20 min with 0.25% trypsin (Invitrogen) at 37°C. After washing, neurons were dissociated using a fire-polished Pasteur pipette and resuspended in Neurobasal medium supplemented with 2% B-27, 1% HEPES, 0.25% GlutaMAX, and 0.1% Penicillin-Streptomycin (all Invitrogen). Neurons were counted in a Fuchs-Rosenthal chamber and plated at 1.5K per well in a 12-well plate. Neuronal cultures were maintained in Neurobasal medium supplemented with 2% B-27, 1% HEPES, 0.25% GlutaMAX, and 0.1% Penicillin-Streptomycin (all Invitrogen), at 37°C in a 5% CO humidified incubator.

Autaptic hippocampal cultures were prepared as described previously [91]. Briefly, micro-islands were prepared with a solution containing 0.1 mg/ml poly-D-lysine (sigma), 0.7 mg/ml rat tail collagen (BD Biosciences) and 10 mM acetic acid (Sigma) applied with a custom-made rubber stamp (dot diameter 250 μm). Next, rat astrocytes were plated at 6-8K per well in pre-warmed DMEM (Invitrogen), supplemented with 10% FCS, 1% Penicillin-Streptomycin and 1% nonessential amino acids (All Gibco).

For rescue experiments, Syt1 KO neurons were infected at DIV4 with a synapsin-promoter-driven lentiviral vector expressing either Syt1 9Pro (residues 264–272 replaced with nine proline residues; kindly provided by dr. Edwin Chapman, Howard Hughes Medical Institute, Madison, WI, USA), Syt1 D232N, Syt1 4W (M173W, F234W, V304W, I367W), or wild type Syt1. The experimental groups were masked during the experiment. The code was broken after statistical analysis.

### Electrophysiology

Whole-cell voltage-clamp recordings (V_m_ = −70 mV) were performed at room temperature with borosilicate glass pipettes (2–5MΩ) filled with (in mM) 125 K^+^-gluconic acid, 10 NaCl, 4.6 MgCl_2_,4 K2-ATP, 15 creatine phosphate, 1 EGTA, and 10 units/mL phosphocreatine kinase (pH 7.30). External solution contained in mM: 10 HEPES, 10 glucose, 140 NaCl, 2.4 KCl, 4 MgCl_2_ (pH = 7.30, 300 mOsmol). 4mM CaCl_2_ was used externally in all experiments, unless otherwise specified. Inhibitory neurons were identified and excluded based on the decay of postsynaptic currents. Recordings were acquired with a MultiClamp 700B amplifier, Digidata 1440 A, and pCLAMP 10.3 software (Molecular Devices). Only cells with an access resistance < 15MΩ (80% compensated) and leak current of <300 pA were included. EPSCs were elicited by a 0.5 ms depolarization to 30 mV.

Hypertonic sucrose stimulation was performed as described previously [7]. Briefly, gravity infused external solution was alternated with 7 s of perfusion with hypertonic solution by rapidly switching between barrels within a custom-made tubing system (FSS standard polyamine coated fused silica capillary tubing, ID 430 μm, OD550 μm) attached to a perfusion Fast-Step delivery system (SF-77B, Warner instruments corporation) and directed at the neuron. Solution flow was controlled with an Exadrop precision flow rate regulator (B Braun). Multiple sucrose solutions with various concentrations were applied to the same cell, taking a 1–2 min rest period in between solutions to accommodate complete recovery of RRP size. In between protocols, a constant flow of external solution was applied to the cells. Multiple sucrose solutions with various concentrations were applied to the same cell, taking a 1–2 min rest period, >3min in case of post-tetanic potentiation protocols. The order of sucrose solutions was alternated between neurons to avoid systematic errors due to possible rundown of RRP size after multiple applications.

For experiments including the cell permeable Ca^2+^ chelator BAPTA-AM, after recording of the first evoked response, cells were incubated for 10 min with 20μM BAPTA-AM (Sigma) in bath and external solution was exchanged for 0mM CaCl_2_. A decrease in spontaneous release during incubation was used as a positive control. For PDBu experiments, sucrose applications were performed as usual, after which neurons were incubated with 1 μM PDBu (Calbiochem), and sucrose applications were repeated.

### Immunocytochemistry

Hippocampal neurons were fixed with 3.7% formaldehyde (Electron Microscopy Sciences) after two weeks in culture. After washing with PBS, cells were permeated with 0.5% Triton X-100 for 5 min and incubated in 2% normal goat serum/0.1% Triton X-100 for 30 min to block aspecific binding. Cells were incubated for 1 h at room temperature with primary antibodies directed against MAP2 and vGlut1 to visualize dendrite morphology and synapses. The following antibodies were used: polyclonal chicken anti-MAP2 (1:10 000, Abcam), polyclonal rabbit vGlut1 (1:500, SySy), polyclonal guinea pig vGlut1 (1:5000, Millipore), monoclonal mouse VAMP2 (1:1000, SySy), polyclonal rabbit Synaptotagmin-1 (1:1000; W855; a gift from T. C. Südhof, Stanford, CA), or polyclonal chicken GFP (1:2000, AVES). After washing with PBS, cells were incubated for 1 h at room temperature with second antibodies conjugated to Alexa dyes (1:1000, Molecular Probes) and washed again. Coverslips were mounted with DABCO-Mowiol (Invitrogen) and imaged with a confocal LSM510 microscope (Carl Zeiss) using a 40× oil immersion objective with 0.7× zoom at 1024 × 1024 pixels. Neuronal morphology was analyzed using a published automated image analysis routine [92], and ImageJ.

### Electron microscopy

Autaptic hippocampal neuron cultures of WT and Syt-1 KO mice (E18) grown on glass cover slips were fixed (DIV14) for 90 minutes at room temperature with 2.5% glutaraldehyde in 0.1 M cacodylate buffer (pH 7.4). After fixation, cells were washed three times for 5 minutes with 0.1 M cacodylate buffer(pH 7.4), post-fixed for 1 hour at room temperature with 1% OsO_4_/1% KRu(CN)_6_. After dehydration through a series of increasing ethanol concentrations, cells were embedded in Epon and polymerized for 48 hours at 60°C. After polymerization of the Epon, the coverslip was removed by alternately dipping it in hot water and liquid nitrogen. Cells of interest were selected by observing the flat Epon-embedded cell monolayer under the light microscope and mounted on pre-polymerized Epon blocks for thin sectioning. Ultrathin sections (80 nm) were cut parallel to the cell monolayer, collected on single-slot, formvar-coated copper grids, and stained in uranyl acetate and lead citrate using a LEICA EM AC20 stainer. Synapses were randomly selected at low magnification using a JEOL 1010 electron microscope. For each condition, the number of docked synaptic vesicles, total synaptic vesicle number, postsynaptic density and active zone length were measured on digital images taken at 80,000-fold magnification using analySIS software (Soft Imaging System). The observer was blinded for the genotype. For all morphological analyses, we selected only clearly recognizable synapses with intact synaptic plasma membranes with a recognizable pre- and postsynaptic area and clearly defined synaptic vesicle membranes. Synaptic vesicles were defined as docked if there was no distance visible between the synaptic vesicle membrane and the active zone membrane. The active zone membrane was recognized as a specialized part of the presynaptic plasma membrane that contained a clear density opposed to the postsynaptic density and docked synaptic vesicles. Cells were cultured from six different WT and seven different Syt-1 KO mice. Approximately 25 synapses were analyzed per culture stemming from one animal.

### Data Analysis

Offline analysis was performed using custom-written software routines in Matlab R2018b (Mathworks). Software routines for analysis of mEPSCs and electrically evoked release is available at https://github.com/vhuson/viewEPSC [93], software for analysis of HS evoked release has been made available previously ([7]; https://doi.org/10.7554/eLife.05531.031). In all figures, stimulation artefacts have been removed. For evoked release, total charge was calculated by integrating the current from the end of the stimulation until the start of the next pulse. HS-induced responses were fitted with a minimal vesicle state model as described previously [7]. This method corrects dynamically for ongoing priming opposed to the use of fixed priming rates in other methods. Furthermore, it provided direct estimates of fusion and priming rates, RRP size, and changes in energy barrier. Parameters describing the kinetics of HS responses were given in supplementary figures for completeness. HS integral was obtained by integrating the full 7s fitted trace. Rise time and time-to-peak were calculated using the minimum of the fitted trace. The delay of HS onset parameter is part of the fitting procedure. Responses to HS concentrations below 500mM were fitted simultaneously with a 500mM response from the same cell, to prevent underestimation of the RRP and overestimation of release rates. The release rate constant during spontaneous release was obtained by dividing the mEPSC frequency by the number of vesicles in the RRP. The latter was calculated by dividing the RRP charge by the average mEPSC charge. Ca^2+^-dependent release rates were simulated using the previously described allosteric model [1]. An RRP size of 15,000 vesicles was chosen based on 500mM HS responses in Syt1 WT. The rate constant *l*_+_ (2.667 × 10^−4^ s^−1^) was set to match spontaneous release in Syt1 WT at 0mM extracellular Ca^2+^ and 20μM BAPTA-AM. All other parameters (*f*: 31.3; *k*_*on*_: 1 × 10^8^ M^−1^ s^−1^; *k*_*off*_: 4,000 s^−1^; cooperativity factor, *b*: 0.5) were set as previously described [1]. Parameters specific to Syt1 D232N (*l*_*+,new*_: 0.00149; *f*_*D232N*_: 62.6) were adapted from Syt1 WT parameters as described in the results section. Statistical significance was determined using Wilcoxon signed-rank tests and Mann-Whitney *U* tests to compare paired- and independent measurements, respectively, p-values below 0.05 were considered significant. All statistical tests were performed in Matlab (Mathworks).

## Supporting information

Statistics Overview

## Acknowledgements

We thank Desiree Schut, Lisa Laan, and Frank den Oudsten for producing glia feeders and primary culture assistance, Joke Wortel for animal breeding, Frank den Oudsten and Joost Hoetjes for genotyping, Robbert Zalm for cloning and producing viral particles, Jurjen Broeke and Hans Lodder for technical assistance.

---

**Figure 1 Supplement 1.**
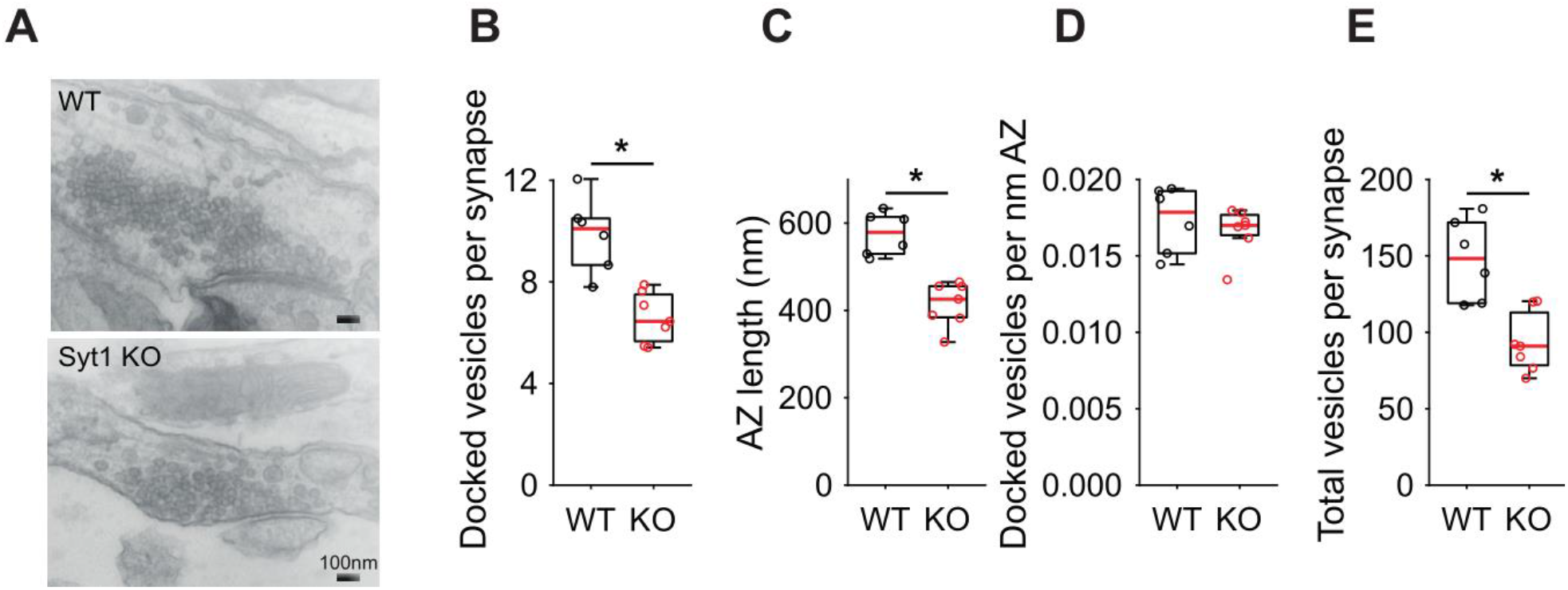
Reduced number of docked vesicles in hippocampal Syt1 KO autaptic synapses. **(A)** Representative examples of electron microscopy images of WT and Syt1 KO synapses. **(B)** Boxplots of docked vesicles per synapses, **(C)** active zone (AZ) length in nm, **(D)** docked vesicles per nm AZ, and **(E)** total number of vesicles per synapse in WT and Syt1 KO. (* *p* < 0.05, Wilcoxon rank sum test).

**Figure 1 Supplement 2.**
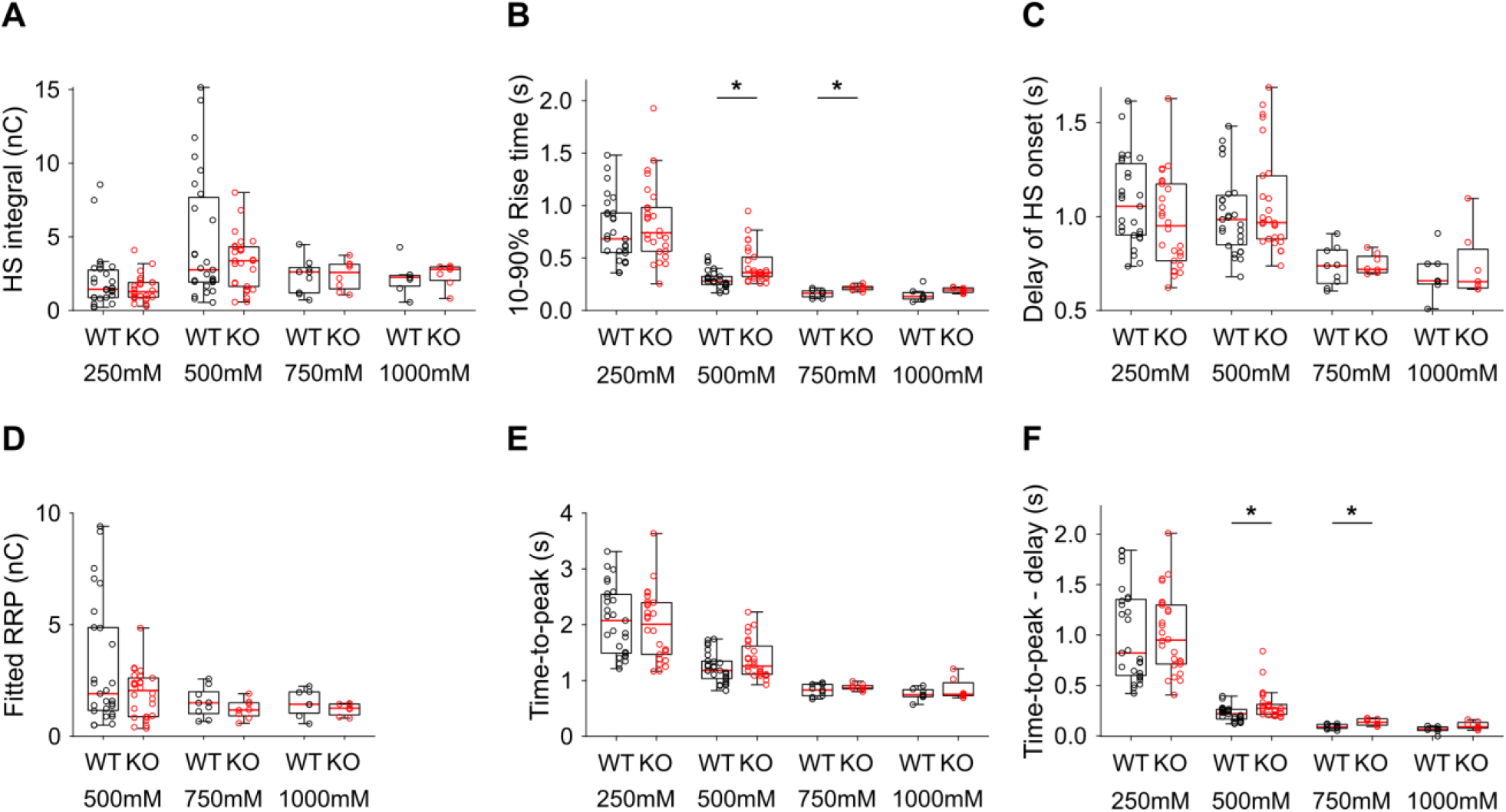
Additional HS parameters from WT and Syt1 KO neurons. **(A)** Integral of current from HS induced release at different concentrations. **(B)** 10-90% rise time of peak HS induced current. **(C)** Time delay after application of HS before onset of release. **(D)** Model derived RRP values from depleting HS stimulations. **(E)** Time from start of HS application till peak HS induced current. **(F)** Time from start of HS application till peak HS induced current, corrected for the delay in release onset after HS application. All panels comparing WT and Syt1 KO (* *p* < 0.05, Wilcoxon rank sum test).

**Figure 2 Supplement 1.**
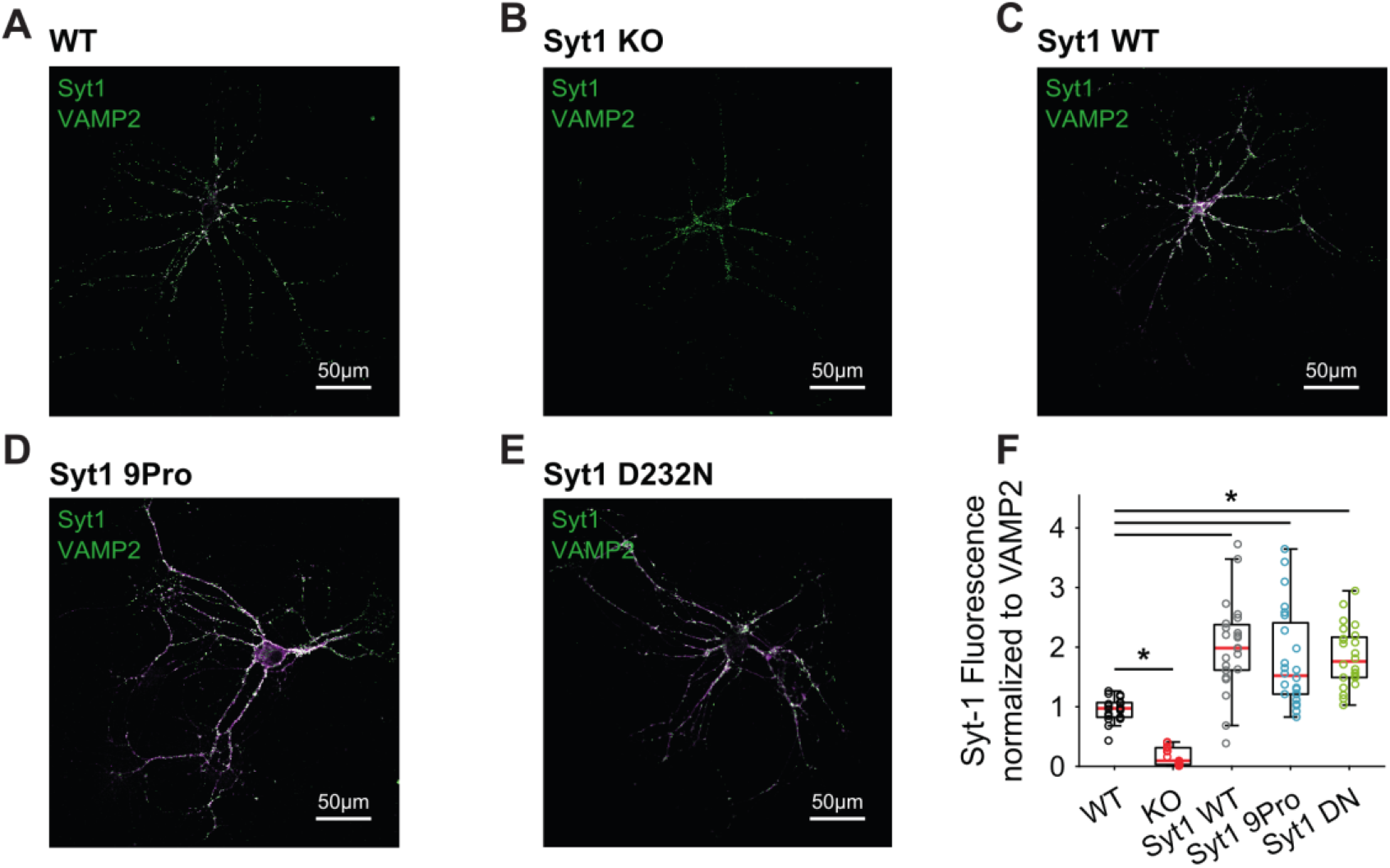
Synaptic expression of Syt1 WT and mutant rescue constructs exceeds endogenous levels. Representative examples of **(A)** WT, **(B)** Syt1 KO, **(C)** Syt1 WT expressing, **(D)** Syt1 9Pro expressing, and **(E)** Syt1 D232N expressing autapses stained for Syt1 (magenta) and the synaptic marker VAMP2 (green). Boxplots of Syt1 fluorescent intensity, normalized to VAMP2 intensity. (* *p* < 0.05, Wilcoxon rank sum test).

**Figure 2 Supplement 2.**
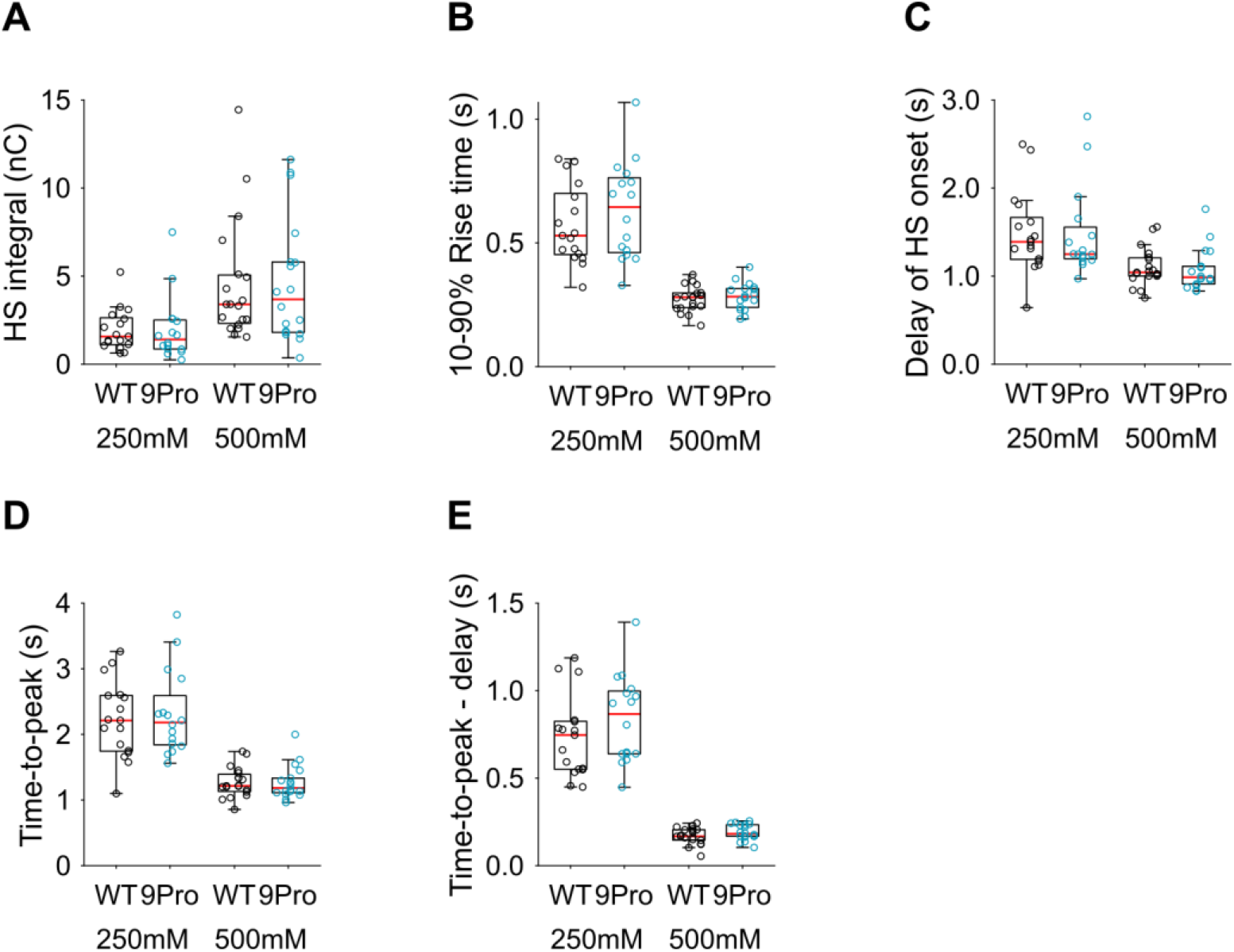
Additional HS parameters from Syt1 WT and Syt1 9Pro expressing neurons. **(A)** Integral of current from HS induced release at different concentrations. **(B)**10-90% rise time of peak HS induced current. **(C)** Time delay after application of HS before onset of release. **(D)** Time from start of HS application till peak HS induced current. **(E)** Time from start of HS application till peak HS induced current, corrected for the delay in release onset after HS application. All panels comparing Syt1 WT and Syt1 9Pro (* *p* < 0.05, Wilcoxon rank sum test).

**Figure 3 Supplement 1.**
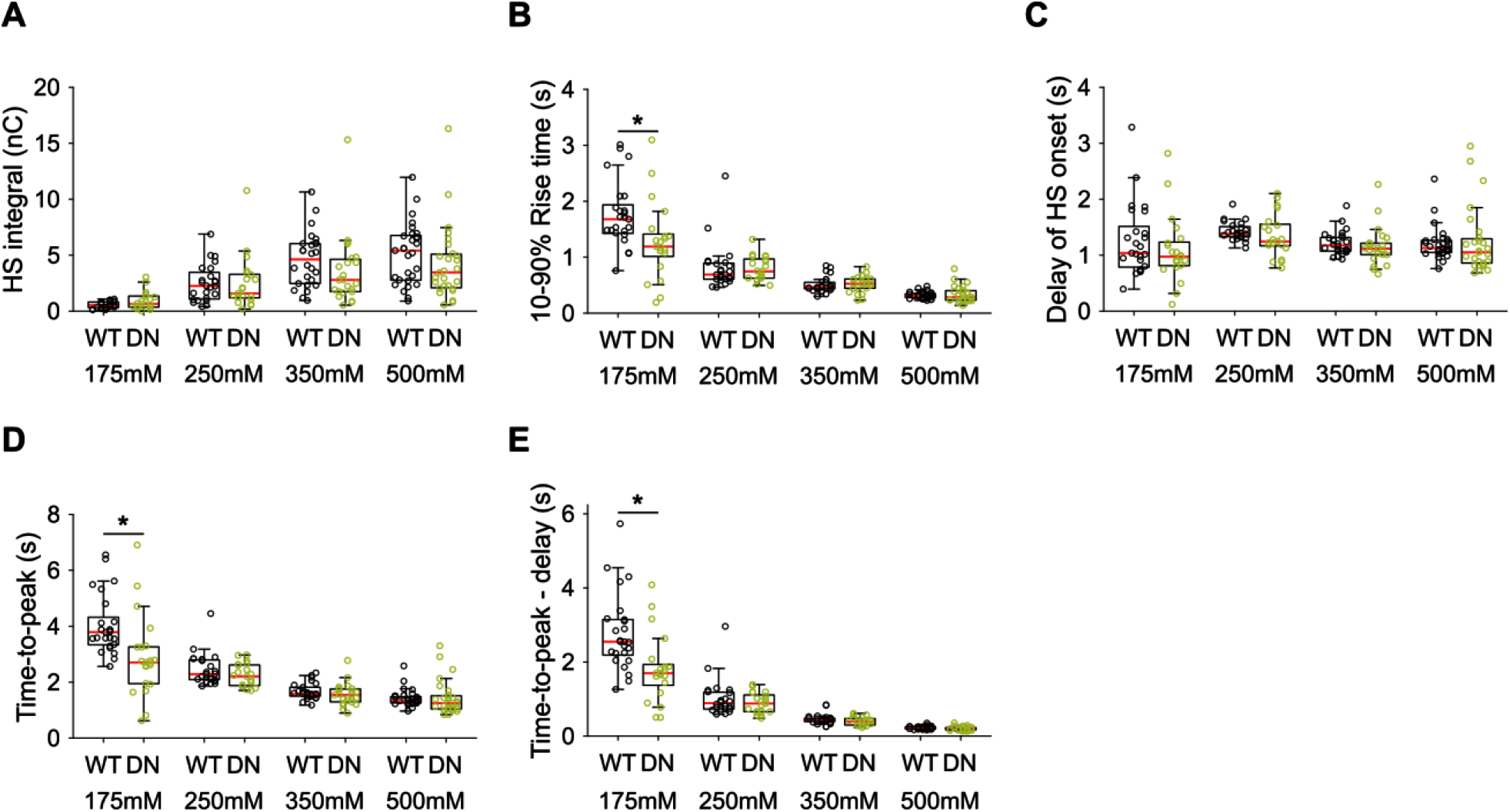
Additional HS parameters from Syt1 WT and Syt1 D232N expressing neurons. **(A)** Integral of current from HS induced release at different concentrations. **(B)** 10-90% rise time of peak HS induced current. **(C)** Time delay after application of HS before onset of release. **(D)** Time from start of HS application till peak HS induced current. **(E)** Time from start of HS application till peak HS induced current, corrected for the delay in release onset after HS application. All panels comparing Syt1 WT and Syt1 D232N (* *p* < 0.05, Wilcoxon rank sum test).

**Figure 3 Supplement 2.**
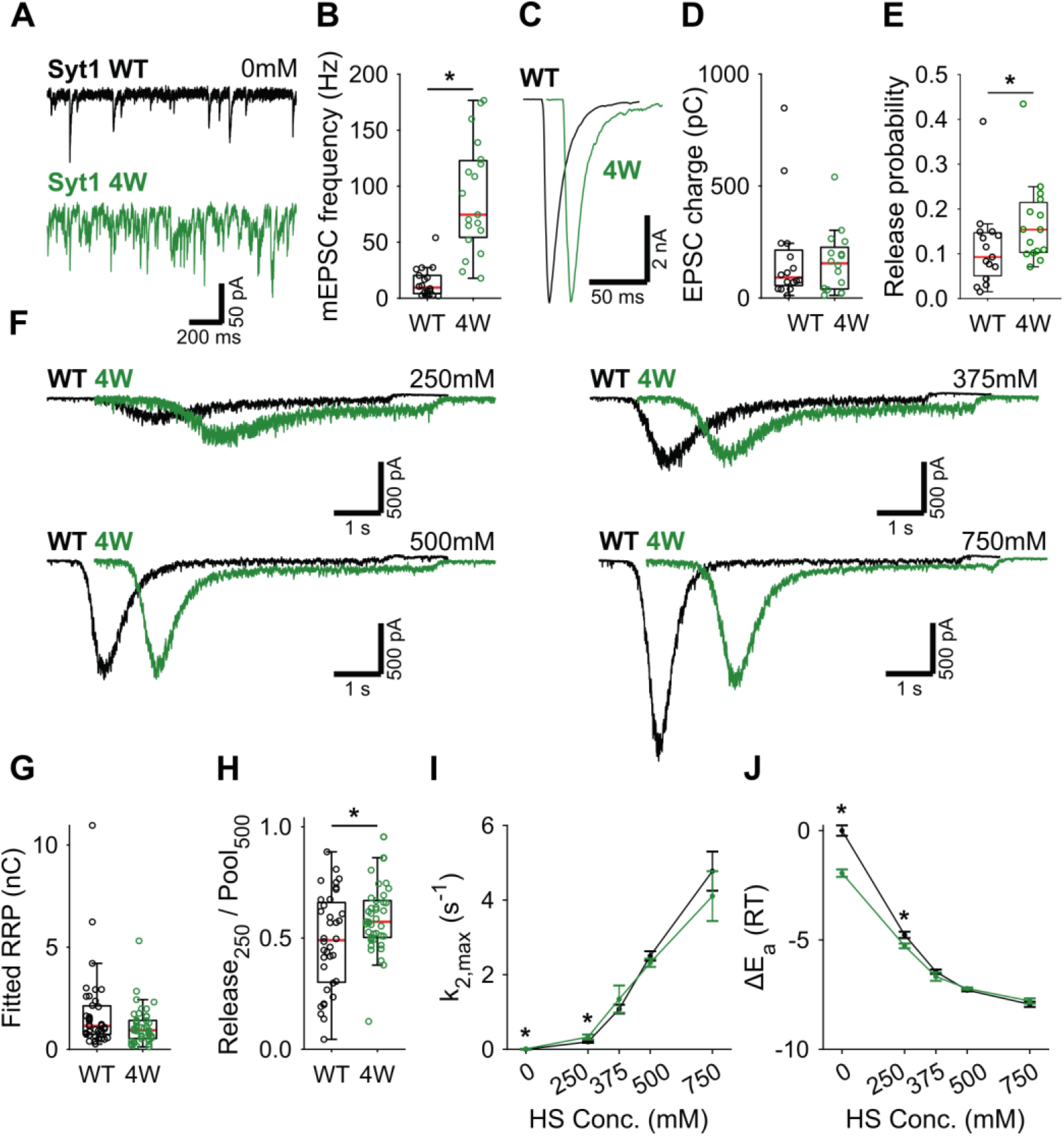
Syt1 4W mutation increases HS release rates only at low concentration. **(A)** Representative traces of spontaneous release (0mM HS), and **(B)** boxplot of spontaneous frequency in Syt1 WT and Syt1 4W expressing synapses. **(C)** Representative traces of AP evoked release in Syt1 WT and Syt1 4W, overlaid with 20ms offset, and **(D)** boxplots of charge transferred during the first evoked EPSC and **(E)** release probability calculated by dividing the EPSC charge by the HS derived RRP charge **(G)**. **(F)** Representative traces of HS induced release in Syt1 WT and Syt1 4W, overlaid with 1s offset, at 250mM, 375mM, 500mM, and 750mM HS, and boxplots of **(G)** RRP charge estimated from 500mM HS, **(H)** depleted RRP fraction at 250mM HS in Syt1 WT and Syt1 4W. **(I)** Plots (mean ± S.E.M.) of maximal HS release rates, and **(J)** change in the fusion energy barrier at different HS concentrations for Syt1 WT and Syt1 4W. (* *p* < 0.05, Wilcoxon rank sum test).

**Figure 3 Supplement 3.**
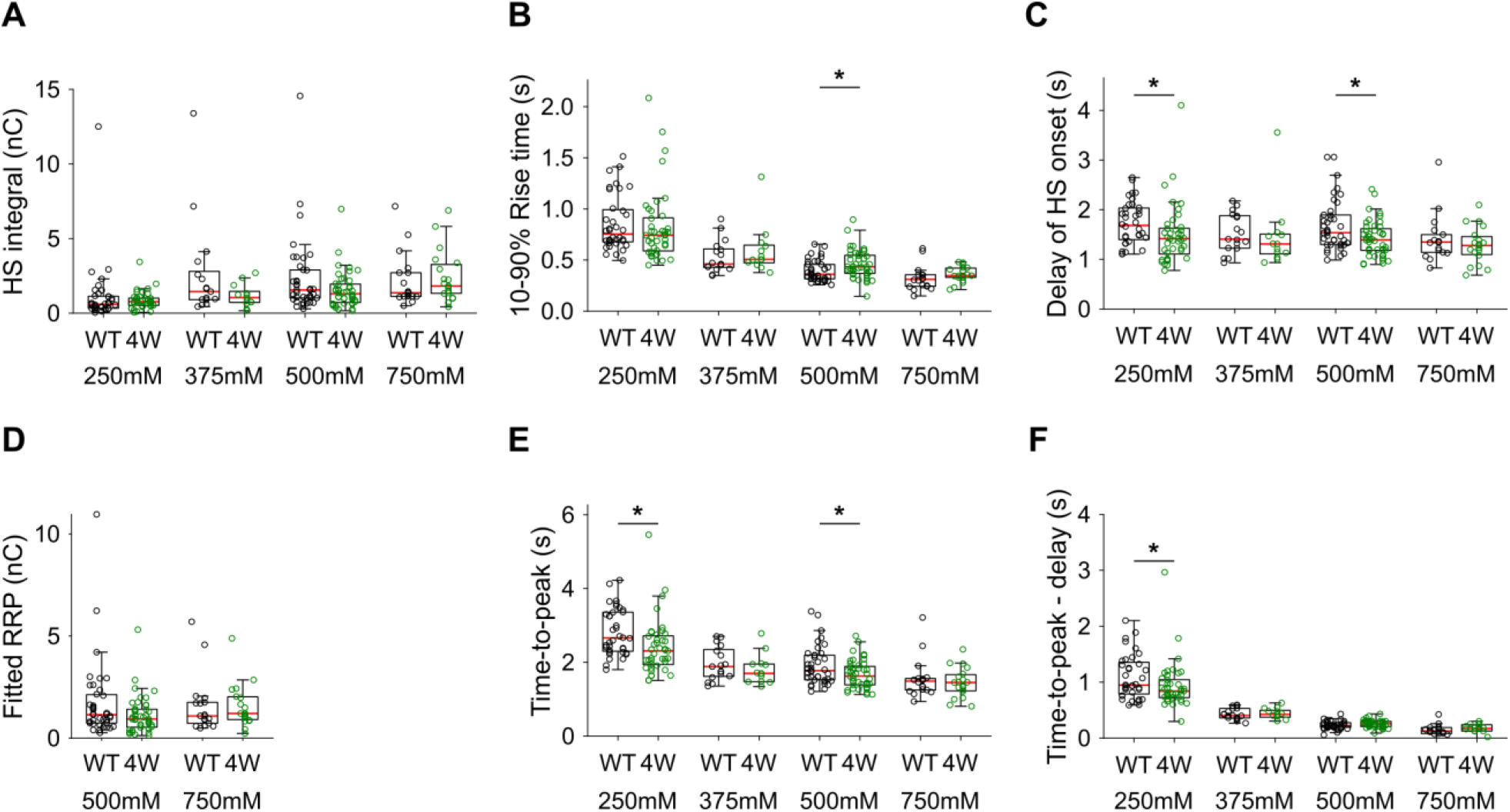
Additional HS parameters from Syt1 WT and Syt1 4W expressing neurons. **(A)** Integral of current from HS induced release at different concentrations. **(B)** 10-90% rise time of peak HS induced current. **(C)** Time delay after application of HS before onset of release. **(D)** Model derived RRP values from depleting HS stimulations. **(E)** Time from start of HS application till peak HS induced current. **(F)** Time from start of HS application till peak HS induced current, corrected for the delay in release onset after HS application. All panels comparing Syt1 WT and Syt1 4W (* *p* < 0.05, Wilcoxon rank sum test).

**Figure 4 Supplement 1.**
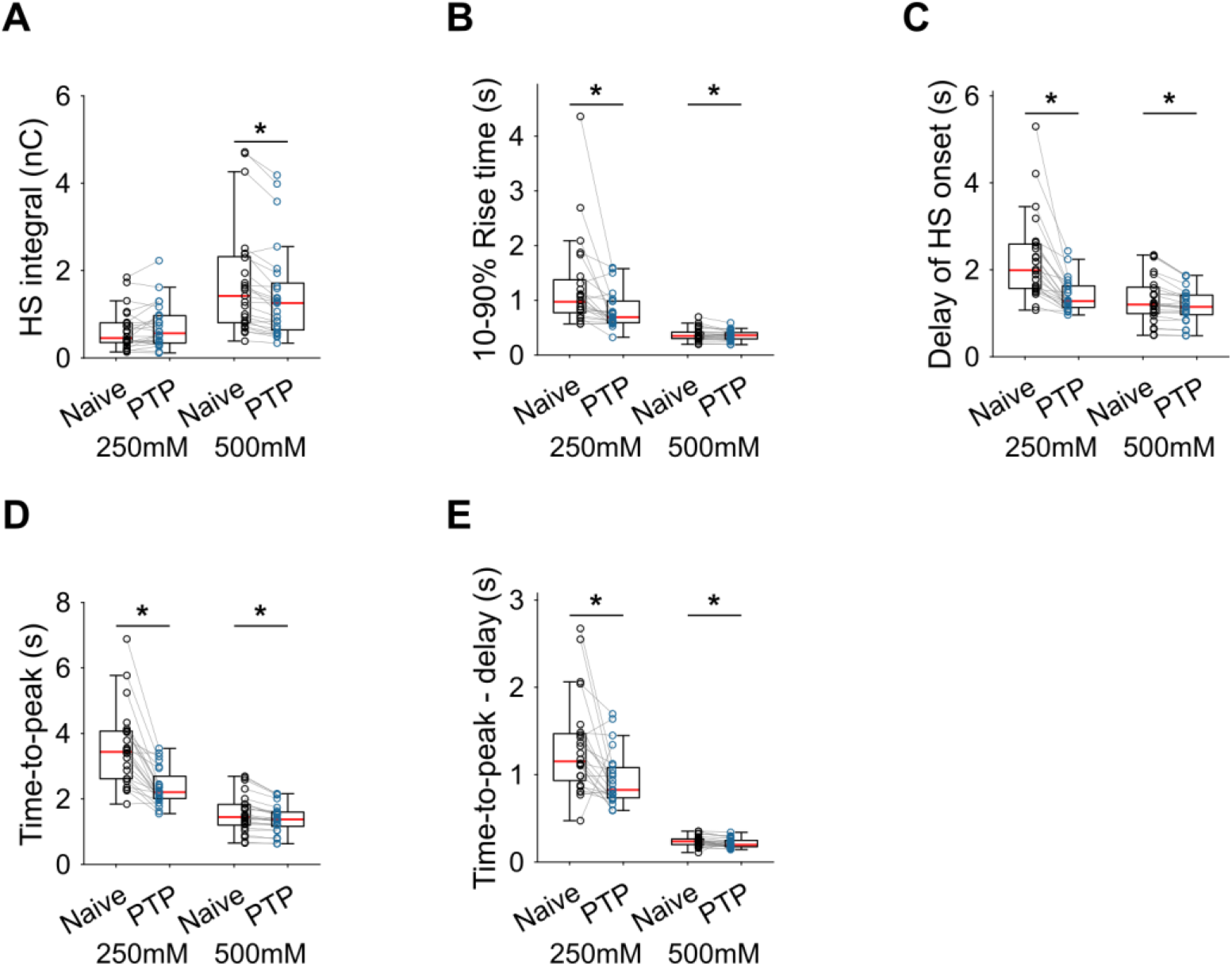
Additional HS parameters before and after PTP in WT neurons. **(A)** Integral of current from HS induced release at different concentrations. **(B)** 10-90% rise time of peak HS induced current. **(C)** Time delay after application of HS before onset of release. **(D)** Time from start of HS application till peak HS induced current. **(E)** Time from start of HS application till peak HS induced current, corrected for the delay in release onset after HS application. All panels comparing before (Naive) and after PTP in WT (* *p* < 0.05, Wilcoxon signed-rank test).

**Figure 5 Supplement 1.**
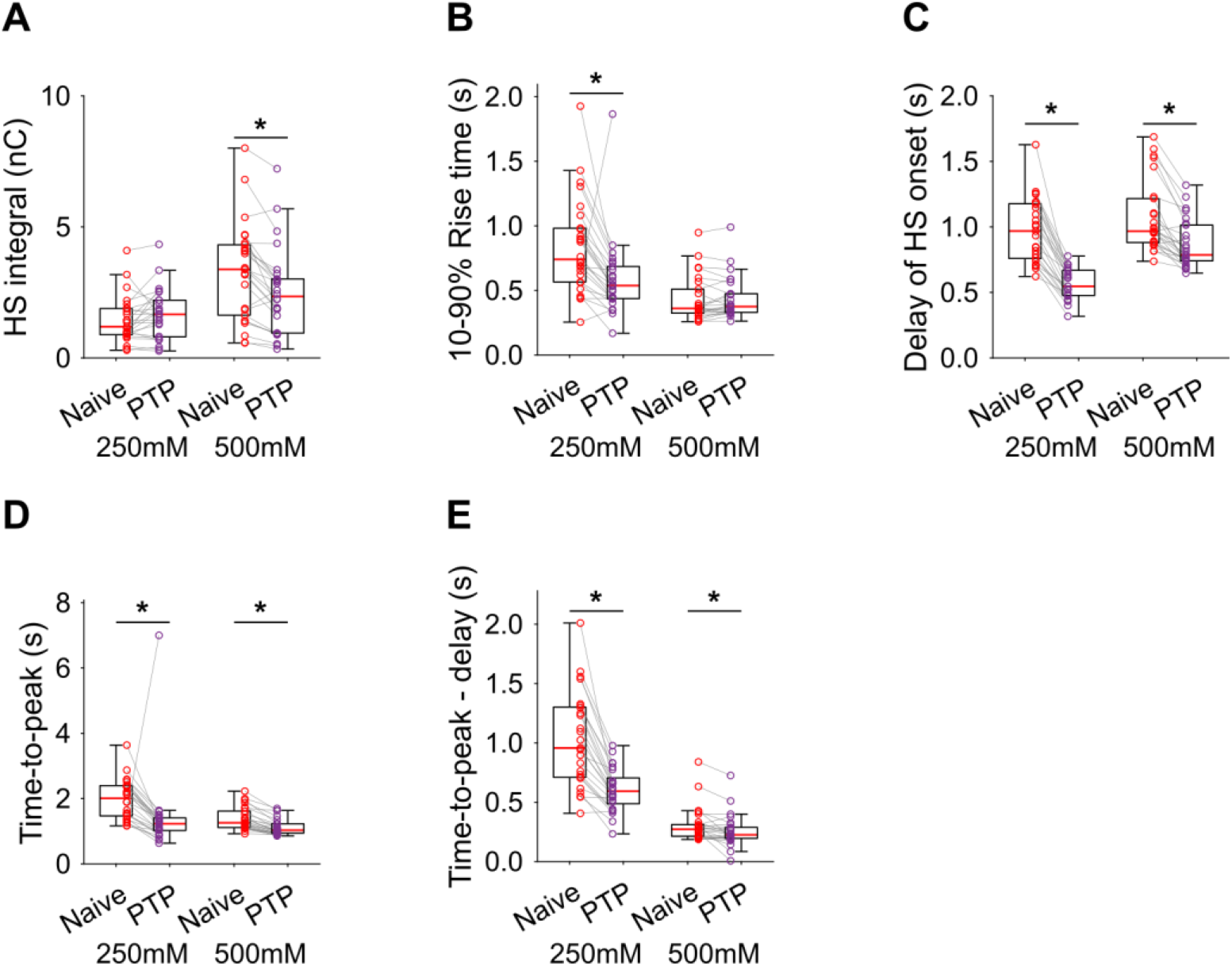
Additional HS parameters before and after PTP in Syt1 KO neurons. **(A)** Integral of current from HS induced release at different concentrations. **(B)** 10-90% rise time of peak HS induced current. **(C)** Time delay after application of HS before onset of release. **(D)** Time from start of HS application till peak HS induced current. **(E)** Time from start of HS application till peak HS induced current, corrected for the delay in release onset after HS application. All panels comparing before (Naive) and after PTP in Syt1 KO (* *p* < 0.05, Wilcoxon signed-rank test).

**Figure 6 Supplement 1.**
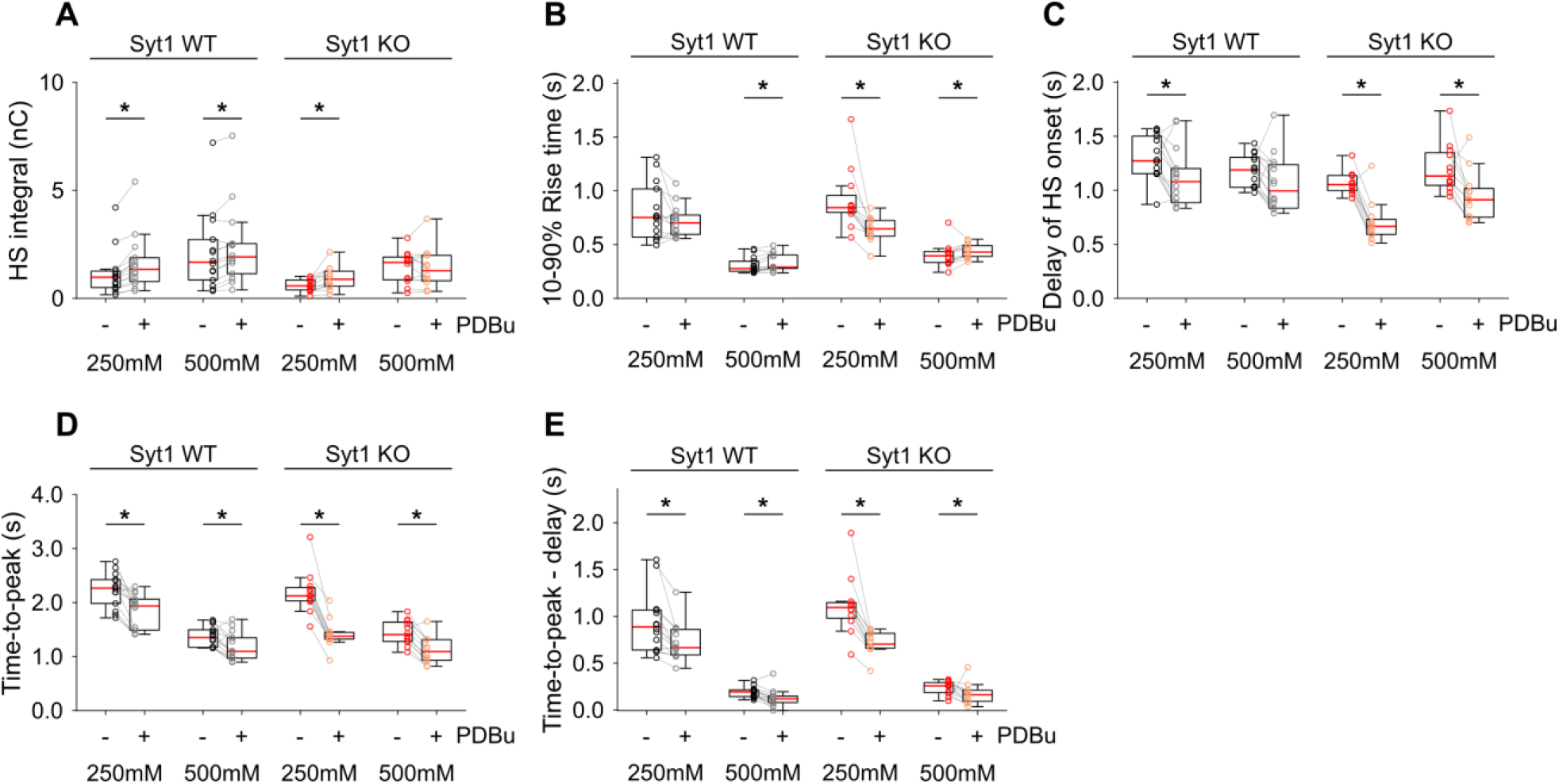
Additional HS parameters before and after PDBu in Syt1 WT and KO neurons. **(A)** Integral of current from HS induced release at different concentrations. **(B)** 10-90% rise time of peak HS induced current. **(C)** Time delay after application of HS before onset of release. **(D)** Time from start of HS application till peak HS induced current. **(E)** Time from start of HS application till peak HS induced current, corrected for the delay in release onset after HS application. All panels comparing before (−) and after (+) PDBu (* *p* < 0.05, Wilcoxon signed-rank test).

**Figure 7 Supplement 1.**
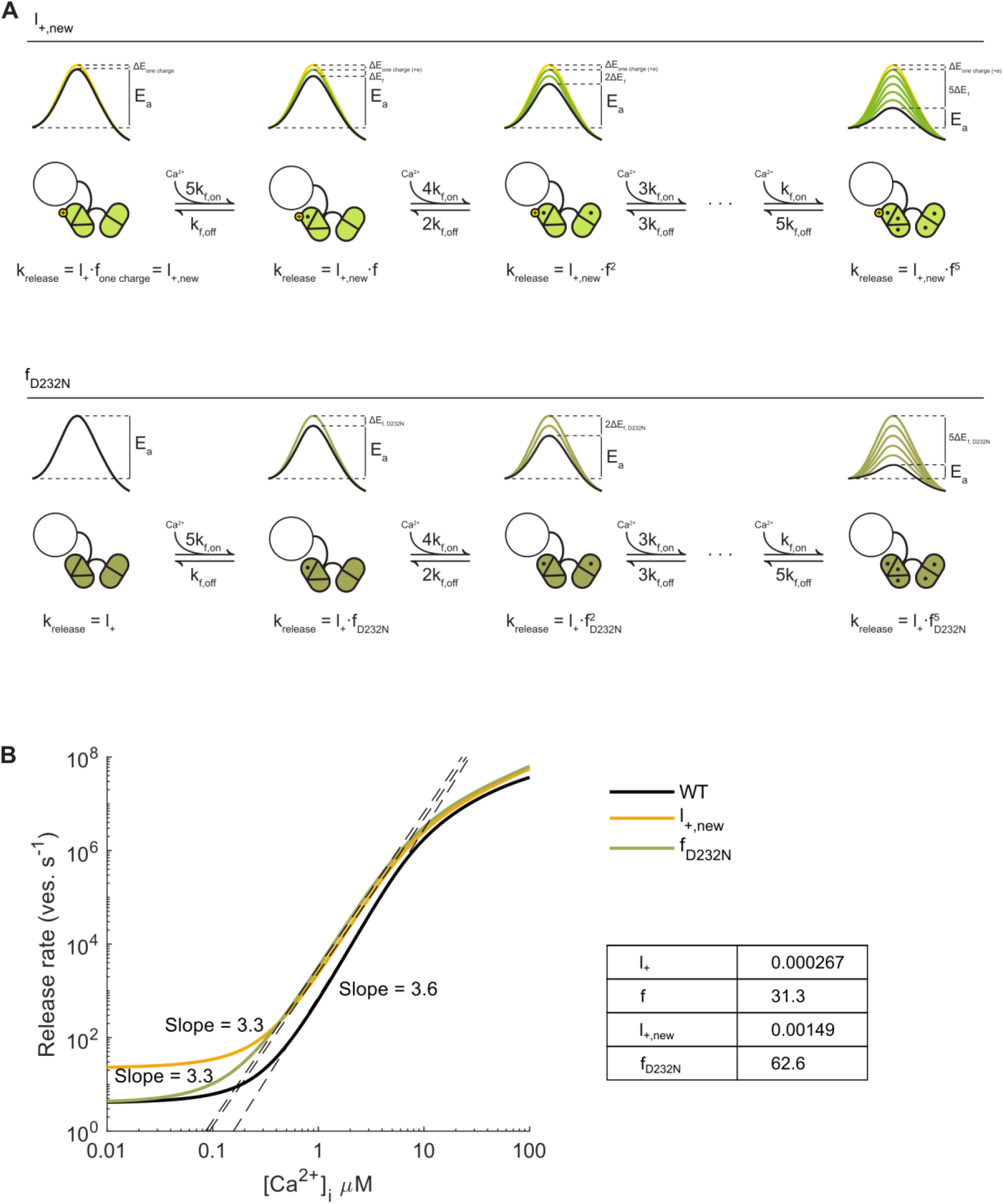
Syt1 D232N as a permanently activated sensor acting on the energy barrier. **(A)** Two scenarios for increased Ca^2+^ sensitivity in Syt1 D232N: 1) Increased positive charge in Syt1 D232N lowers the ground state energy barrier, thereby increasing the basal fusion rate constant (*l*_+_), with half the efficacy of a single Ca^2+^ ion, to 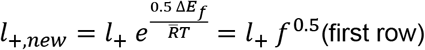. 2) Twofold increase in Ca^2+^-dependent SNARE binding in Syt1 D232N, increases the effect of Ca^2+^ binding on the fusion rate two-fold (*f*_*D232N*_ = 2*f*) (second row). **(B)** Simulations with the allosteric model [1], reveal that both increased positive charge (*l*_+,*new*_) and increased SNARE binding (*f*_*D232N*_) lead to higher release rates, though only for *l*_+,*new*_ was Ca^2+^-independent release also increased. Linear fits of the steepest part of the curves indicate no clear change in Ca^2+^-cooperativity compared to WT.

